# Understanding *Wolbachia* acquisition and co-divergence of hosts and their associated bacteria: *Wolbachia* infection in the *Chorthippus parallelus* hybrid zone

**DOI:** 10.1101/044784

**Authors:** Paloma Martinez-Rodriguez, Francisca Arroyo-Yebras, Jose Luis Bella

## Abstract

*Wolbachia* is one of the best known bacterial endosymbionts affecting insects and nematodes. It is estimated that it infects 40% of insect species, so epidemiologically it may be considered a pandemic species. However, the mechanisms by which it is acquired from other species (horizontal transmission) or by which it coevolves with its hosts as a result of vertical transmission across generations are not known in detail. In fact, there are few systems in which the codivergence between host and bacterium has been described.

This study goes in deep in the *Wolbachia* infection in the grasshopper *Chorthippus parallelus*. This well-known system allows us to investigate the mechanism of acquisition of various *Wolbachia* strains in a new host, and the bacterial genomic changes during bacterial-host codivergence: We describe the genetic diversity of *Wolbachia* strains infecting both subspecies of *C. parallelus* and analyse their phylogenetic relationship. We also show the emergence of new bacterial alleles resulting from recombination events in *Wolbachia* infecting hybrid hosts. Our data suggest that F strains detected in this grasshopper have co-diverged with its host, *versus* a more recent horizontal transmission of B strains. According with this, we discuss the potential role of *Wolbachia* in the dynamics of the grasshopper hybrid zone and in the divergence of the two grasshopper subspecies since the origin of their hybrid zone.

## Introduction

*Wolbachia* is one of the most widely distributed endosymbiotic bacteria, infecting about 40% of insect species. At least, 8 bacterial supergroups have been described (Zug & Hammerstein 2012, but see Gerth et al. 2014). Vertical (from females to offspring) and horizontal (across species) transmission are the two main mechanisms to explain *Wolbachia* expansion. However, the way in which the two main modes of transmission have combined during the evolutionary history of *Wolbachia* and its hosts it is not well understood (Kremer & Huigens 2011; Werren et al., 2008). On the one hand, horizontal transmission has been proposed as an essential mechanism to explain the current distribution of *Wolbachia* across species. Actually, horizontal transmission and infection loss could explain the observed phylogenetic incongruence between *Wolbachia* and its hosts or the appearance of the same *Wolbachia* strain in distantly related host species (Baudry *et al*. 2003; Keller *et al*. 2004; Martins *et al*. 2012; Raychoudhury *et al*. 2009; Shoemaker *et al*. 2003; Yun *et al*. 2011). On the other hand, vertical transmission is the predominant mode of transmission (Moran *et al*. 2008; Saridaki & Bourtzis 2010). Due to that, coevolution between *Wolbachia* and their host should be common (but it has been rarely described) (Raychoudhury *et al*. 2009; but see Bordenstein *et al*. 2009 and Gerth *et al*. 2014). Here, the *C. parallelus* hybrid zone was used to investigate this infrequently reported process due to the knowledge about the evolutionary history of this species.

The hybrid zone formed by the meadow grasshopper *Chorthippus parallelus* is considered an example of secondary contact after allopatric differentiation (Bella *et al*. 2007; Hewitt 1993; Shuker *et al*. 2005). After the last ice age, *C. p. parallelus* and *C. p. erythropus* met at the geographical barrier of the Pyrenees, giving rise to the hybrid zone that exists to this day. Currently, the hybrid zone along the valleys of Tena (Spain) and d’Ossau (France) extends over more than 40 km, where a gradient of phenotypic and genotypic characters have been found between pure populations, located at the ends of the hybrid zone (Hewitt 2001; Hewitt 2011; Shucker *et al*. 2005).

B and F *Wolbachia* supergroups infect *C. parallelus* (Dillon et al., 2008; Martínez *et al*. 2009; Zabal-Aguirre *et al*. 2010). Previous data provide evidence of different patterns of infection and coinfection by the two bacterial supergroups in pure and hybrid populations throughout the Iberian Peninsula, the Pyrenees and the rest of Europe, based on infection frequencies (Bella *et al*. 2010; Martinez-Rodriguez 2013; Zabal-Aguirre *et al*. 2010). It is noteworthy that these bacterial biogeographical patterns clearly delineate the current distribution of pure and hybrid grasshoppers.

Experimental crosses in the field with pure and hybrid individuals of *C. parallelus* show that *Wolbachia* causes cytoplasmic incompatibility in crosses between infected and uninfected individuals (unidirectional incompatibility) and in those between individuals infected with different bacterial lineages (bidirectional cytoplasmic incompatibilities), as indicated by the significant reduction in the number of offspring of the affected crosses. *Wolbachia* also increases the fecundity of infected females (Zabal-Aguirre *et al*. 2014). In addition, the bacterium induces certain cytogenetic effects in this grasshopper, this affecting the proportion of abnormal spermatids and the chiasmata frequency (Sarasa *et al*. 2013). The existence of CI and other mentioned effects suggest that *Wolbachia* infection could influence the dynamics of the *Chorthippus* hybrid zone, reinforcing the reproductive barrier between them. Actually, several theoretical studies support this fact: For example, recently Telschow *et al*. (2014) report that nuclear incompatibilities (according with Dobzhansky Muller model) and cytoplasmic incompatibilities could act synergistically in order to keep the existence of genetic diversity after secondary contact. However, more studies are required to understand the underlying processes in this particular case.

In this study and based on the multilocus system typing (MLST system) proposed by Baldo *et al*. (2006b), we (1) analyse the phylogenetic relationship between *Wolbachia* strains infecting host populations and (2) the current distribution of *Wolbachia* infection in pure and hybrid populations of this grasshopper inside and outside its hybrid zone, including populations outside the Iberian peninsula. This also serves to propose the possible influence of ancestral F *Wolbachia* in the very origin of this hybrid zone. Besides we (3) describe the greater genetic variability in *Wolbachia* strains infecting grasshopper hybrid populations *vs*. pure subspecies populations, which suggests close endosymbiont/host-genotype interactions and provides evidence of coupled evolution between both genomes. Finally, we infer (4) the modes of acquisition of *Wolbachia* in *C. parallelus* and describe (5) how the combination of vertical and horizontal modes of transmission explains current patterns of *Wolbachia* infection in *C. parallelus* and its consequences for the evolutionary history of the host.

## Material and methods

### Field collections

*Wolbachia* infection was analyzed in more than 1780 *Chorthippus parallelus* individuals collected from 21 European locations inside and outside of the hybrid zone in 2008 and 2009, with the exception of Bubion and Epping Forest populations, captured in 2002 and 2004, in the context of a *Wolbachia* infection prevalence experiment in *Chorthippus* (see Martinez-Rodriguez, 2013). The populations are grouped as indicated in Table 1. Complete data collection are indicated in supplementary table 1. Gonads were dissected and fixed in 100% ethanol.

**Table 1.** Coordinates and designation of sampled populations of *C. parallelus*. (1) Hybrid population according to (Serrano *et al*., 1996). (2) Pure *Chorthippus parallelus* population, with some particular cytogenetics markers (see Bella *et al*. 2007). (3) Hybrid population in northern Spain (as characterized by Bella *et al*. 2007). (4) Hybrid population, according to (Flanagan *et al*. 1999) between *C. parallelus parallelus* and additional (5) Italian subspecies.

| Population nomenclature (in figures) | Population | Hybrid-pure status | Latitude | Longitude | Altitude (m) |
| --- | --- | --- | --- | --- | --- |
| **Pyrenees-Hybrid Zone** |  |  |  |  |  |
| HZ France | Arudy (France): ARU | Pure Cpp population | 43° 06' 01" N | 0° 26' 38" W | 411 |
|  | Gabas (France): GAB | Pure Cpp population | 42° 53' 60" N | 0° 25' 60" W | 1020 |
|  | L'Hermine (France): HER | Hybrid population | 42° 51' 46.8" N | 0° 23' 30.4" W | 1209 |
|  | Soques (France): SOQ | Hybrid population | 42° 20' 08" N | 0° 23' 52" W | 1396 |
| Tourmont | Cabaña Tourmont (France): TOU | Hybrid population | 42° 49' 11" N | 0° 24' 21" W | 1625 |
| Portalet | Portalet (Spain): POR/PORCRU | Hybrid population | 42° 48' 03" N | 0° 24' 54" W | 1780 |
| CM | Corral de Mulas (Spain): CM | Hybrid population | 42° 47' 09.4" N | 0° 23' 34.4" W | 1569 |
| Sallent | Sallent de Gállego (Spain): SAL | Hybrid population<sup>1</sup> | 42° 45' 57.5" N | 0° 20' 33.9" W | 1343 |
| Escarrilla | Escarrilla (Spain): ESC | Pure Cpe population | 42° 43' 54.1" N | 0° 18' 39.3" W | 1130 |
| **Pyrenees (other)** |  |  |  |  |  |
| South-Pyrenees/Vielha | Puerto del Cantó (Spain): PCAN | Pure Cpe population | 42° 22' 12.9" N | 1°14'11.7" E | 1725 |
|  | Muna (France): MUN | Pure Cpp population | 42° 53' 53" N | 0° 37' 48.8" E | 544 |
|  | Vielha (Spain): VIEL | Pure Cpe population | 42° 40' 25.3" N | 0° 46' 26.5" E | 1393 |
| **Iberian peninsula** |  |  |  |  |  |
| Centre | Navafria (Spain): NAV | Pure Cpe population | 40° 59' 01.95" N | 3° 49' 00.9" W | 1780 |
|  | Becedas (Spain): BEC | Pure Cpe population | 40° 24' 18" N | 5° 38' 17.2" W | 1091 |
| Bubion | Bubion (Spain): BUB | Pure Cpe population | 36° 57' 1.8" N | 3° 21' 22.8" W | 1332 |
| North | Basque Country I (Spain): ALA | Pure Cpe population<sup>3</sup> | 42° 58' 41.4" N | 2° 44' 19.7" W | 625 |
|  | Basque Country II (Spain): URK | Pure Cpe population<sup>3</sup> | 43° 13' 59.1" N | 2° 29' 22.3" W | 211 |
| Europe |  |  |  |  |  |
| Alps | Valdieri (Italy): VAL | Pure Cp population<sup>5</sup> | 44° 12' 19.74" N | 7° 22' 47.76" E | 983 |
|  | Col de L'Arche (France): CLAR | Hybrid population<sup>4</sup> | 44° 25' 34.3" N | 6° 53' 21.6" E | 1942 |
| UK | Epping Forest (England): ING | Pure Cpp population | 51° 39' 36" N | 0° 3' 0" E | 102 |
| Slovenia | Mokronog (Slovenia): SLO | Pure Cpp population | 45° 56' 37.17" N | 15° 8' 55.428" E | 242 m |

### DNA extraction, Wolbachia detection and sequencing

DNA was extracted from whole fixed ovaries and testes, as described in Martínez-Rodríguez *et al*. 2013a and 2013b. *Wolbachia* was detected by PCR amplification of a *Wolbachia 16S rRNA* gene in all sampled individual, using *Wolbachia-*specific primers (Zabal-Aguirre *et al*. 2010), followed by a second, nested PCR amplification using strain-specific primers (Martínez-Rodríguez *et al*. 2013a and 2013b) (Table 2). PCR and Nested-PCR reactions were adjusted to 25 µl: 1X buffer, 2 mM of Mg2Cl, 0.2 mM dNTP, 1.2 µM each primer, 1.25 units of BioTaq DNA polymerase (Bioline) and 100 ng genomic DNA (for the first PCR) or 0,5 µl of previous PCR product in the nested PCR). The reaction was initiated with a cycle of 95° C 30s, followed by 35 cycles of 30s at 95° C, 1 min at 54°C (first PCR) or 69° C (nested-PCR), 1min 30s at 72° C and a final cycle of 10 min at 72° C. A total of 10 µl of each amplification product were electrophoretically separated on 1% agarose gels, which were stained with 0.5 mg/ml ethidium bromide and visualized under UV light (UVIdoc, Uvitec Cambridge).

**Table 2.**
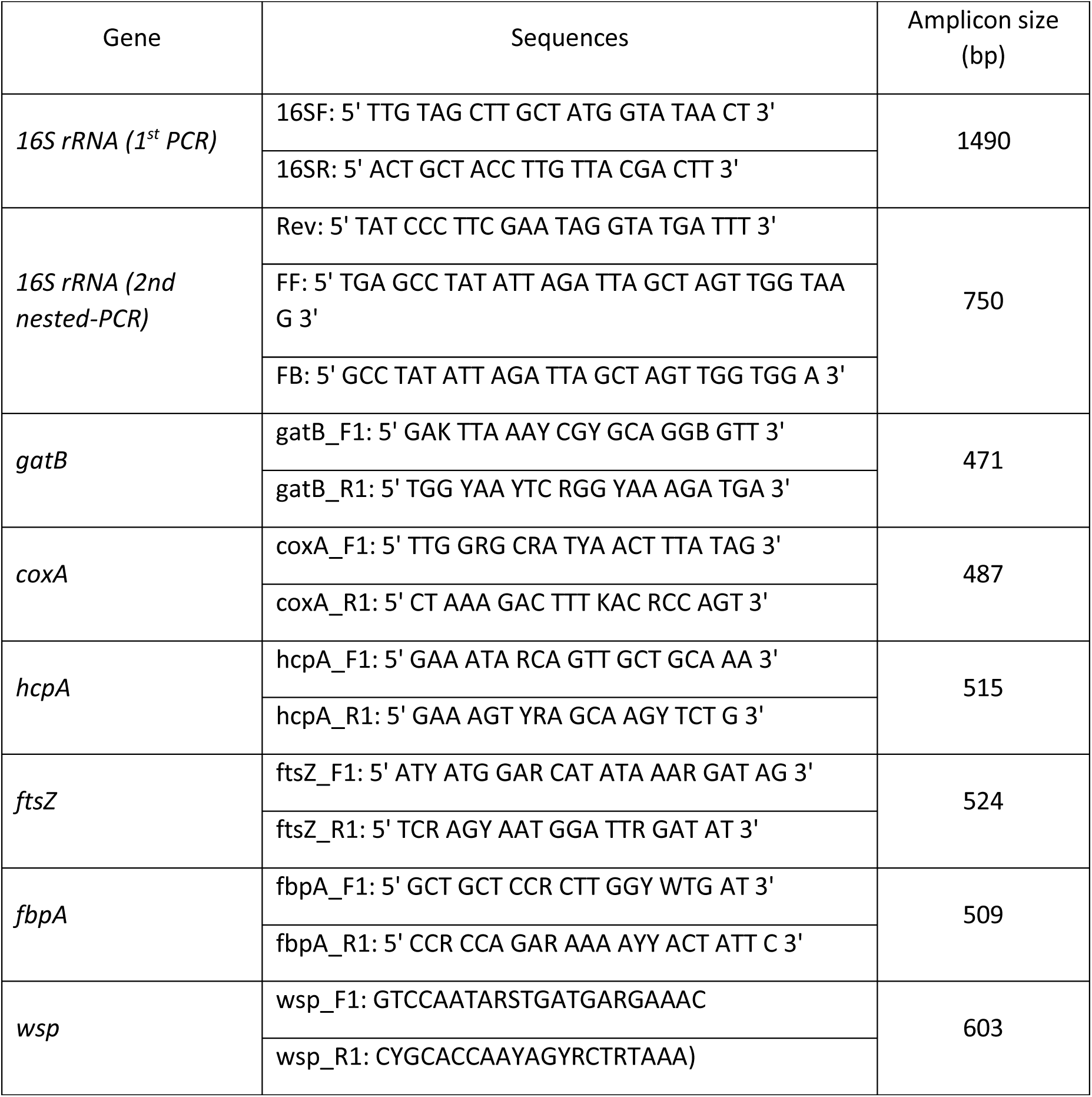
Primer sequences used in the study.

We characterized the *Wolbachia* strains using the MLST and *wsp* (*Wolbachia* surface protein) gene characterisation systems (Baldo *et al*. 2006b). The *gatB, coxA, hcpA, ftsZ, fbpA* and *wsp* genes were amplified in 127 selected singly infected individuals (individuals infected exclusively by F or B supergroup, according with the previous *16S rRNA* gene test, supplementary table 1) while multiple-infected individuals were discarded to avoid ambiguous chromatogram lectures and to reduce the experimental workload. Infection frequencies according with 16S gene test in the different populations are described in Martinez-Rodriguez, 2013 and supplemental table 1). These genes were amplified using previously described methods (Baldo *et al*. 2006b) with slight modifications: PCR reactions were performed in 50 µl volumes containing 2 mM of MgCl_2_, 0.2 mM of dNTP, 30 pmoles of each primer, 1.25 U of *Taq* BIOTAQ™ DNA polymerase (Bioline) and 2 µl of DNA solution (50 ng/µl). The reaction was initiated with a cycle of 95° C 30s, followed by 35 cycles of 30s at 95° C, 1 min at 54°C (*hcpa, gatb, ftsz* and *coxa* genes) or 59° C (*wsp* and *fbpa*), 1min 30s at 72° C and a final cycle of 10 min at 72° C. A total of 10 µl of each amplification product were electrophoretically separated on 2% agarose gels, which were stained and visualized as described above. Amplified genes were purified by ExoSAP-IT (GE Healthcare) and Sanger automatically sequenced (by Stabvida, Portugal). The MLST and *wsp* sequences generated in this study have been deposited in the GenBank database under accession numbers KM078849-KM078883 (see supplementary table 2).

### Sequences analyses

Further studies in this grasshopper confirm *Wolbachia* integrations in the host genome. Current genomic data confirm the absence of integrated sequences of *ftsZ, fbpA* and *wsp* genes. However, some incomplete reads that mapped to *coxA, gatB* and *hcpA* have been detected in uninfected individuals in low coverage (Funkhouser-Jones *et al*., 2015). This forces to be cautious before confirming that the sequences obtained by PCR belong to infecting bacteria and an accurate protocol was developed to ensure this, as indicated below.

Firstly, DNA was extracted from gonad tissue in order to increase the living bacteria/nuclear insertions ratio. Previous studies confirm the massive presence of *Wolbachia* in the grasshopper gonads of infected individuals (Martínez *et al*., 2009). This reduces the probability of Sanger sequencing *Wolbachia* insertions. Secondly, to distinguish the sequences belonging to infecting *Wolbachia* and those sequences integrated into the host nucleus all sequences were compared with the standard sequences to detect possible rearrangements and also translated into protein, in order to detect frameshift mutations, stop codons, and indels. Previous studies in a different grasshopper, *Podisma pedestris*, confirm that most *Wolbachia* insertions show these types of mutations, due to the absence of evolutionary constraints after integration (non-translated sequences) (Martinez-Rodriguez *et al*. unpublished data). Although this reduces the probability of considering an integrated sequence as belonging to “living *Wolbachia*”, we are reminded that it cannot totally discard this possibility. We are taking this in mind when describing and discussing our results, mainly those regarding recombinant and new alleles (see below).

### Phylogenetics analysis

Bayesian likelihood was inferred using a Markov Chain-Monte Carlo variant run in the MrBayes 3.2.1 program (Ronquist & Huelsenbeck 2003). Phylogenies based on single and concatenated MLST genes and *wsp* were reconstructed. JModeltest (Posada 2008) was used to distinguish the appropriate model of evolution, the best likelihood score being chosen on the basis of the AIC criteria (Akaike 1974). The selected models were GTR+I+G for concatenated MLST, *ftsZ* and *gatB*; GTR+G (general time-reversible model, including gamma correction) for *coxA, hcpA* and *wsp*; and HKY+I+G (the Hasegawa, Kishino and Yano model (Hasegawa *et al*., 1985), including gamma and proportion invariant corrections) for the *16S rRNA* gene. Bayesian analysis was carried out for 10^6^ generations with a sample frequency of 100. The first 25% of trees were considered as burn-in and thus discarded. For each locus, the level of nucleotide diversity per site and the number of variable sites or Ka/Ks were estimated using DnaSP software (Librado & Rozas 2009). Alignments of individual and concatenated genes with and without outgroups were screened for significant levels of recombination using RDP4 v4.16 (Martin *et al*. 2010). The analysis involved several tests including GENECONV (Padidam *et al*. 1999), MAXCHI (Maynard Smith 1992) and Chimaera (Posada & Crandall 2001). A Bonferroni correction was applied and significance was concluded for values of p < 0.01.

### Strain characterisation

Following the MLST system (Baldo *et al*. 2006b; Maiden *et al*. 1998), we defined a *Wolbachia* strain or sequence type (ST) as being different on the basis of its unique combination of five alleles. Furthermore, strains sharing at least three alleles were considered to belong to an ST complex, a group of evolutionarily related haplotypes. This analysis was carried out using START2 (Jolley *et al*. 2001). The *wsp* system was employed as a complementary approach for strain characterisation (see Baldo *et al*. 2005, Baldo *et al*. 2006a). Alleles that were detected only once were excluded in the analysis to avoid the miss interpretation of the PCR-associated sequencing errors.

### Inference of bacterial microevolution using multilocus sequence data

We inferred *Wolbachia* microevolution using ClonalFrame to identify the clonal relationships between strains, and to estimate recombination events that have disrupted the clonal inheritance (Didelot & Falush 2007). We performed five separate runs, executing 250,000 MCMC iterations for each, discarding the first 100,000 iterations as burn-in.

### Biogeographical analysis

An AMOVA based on the ST frequencies detected in each population was carried out based on the estimated supergroup frequencies (some data here used from Bella *et al*. 2010 and Zabal-Aguirre *et al*. 2010) and the genetic distance between haplotypes (calculated as the Tamura–Nei distance). Locus-by-locus AMOVA and an exact test of population differentiation were also carried out. In addition, we tested the correlation between genetic and geographical distances with Mantel tests. Geographical distance was estimated using Geographical Distance Matrix Generator v.1.2.3 (http://biodiversityinformatics.amnh.org). All analyses were done using Arlequin 3.11 (Excoffier *et al*. 2005).

## Results

### 1 Wolbachia diversity in C. parallelus

#### 1.1 How many *Wolbachia strains infect C. parallelus*?

To characterize the *Wolbachia* diversity across the hybrid zone, *16S rRNA*, MLST and *wsp* genes of *Wolbachia* were sequenced from host grasshopper individuals collected in several populations, inside and outside the hybrid zone.

The reanalysed phylogenetic tree based on *Wolbachia 16S rRNA* gene sequences confirmed that *C. parallelus* are infected by at least 4 strains belonging to the F supergroup and 2 B supergroup’s strains (Bella *et al*. 2010; Martínez-Rodríguez *et al*. 2013a; Zabal-Aguirre *et al*. 2010) (see supplementary Fig. 1).

In addition, we studied *Wolbachia* supergroups and strains infecting *C. parallelus*, on the basis of the five genes involved in the MLST system, and on the *wsp* gene (Baldo *et al*., 2006b). The analysis of the sequences of the 5 MLST genes distinguish 33 different haplotypes or ST (sequence types, according with Baldo *et al*. 2006b) based on the combination of 5 loci alleles: We detected 5 different alleles of ftsZ gene, 5 alleles of gatB gene, 6 alleles of coxA gene, 5 alleles of fbpA gene, and 10 alleles of hcpA gene (see Fig. 1, 2 and supplemental Figs. S2 to S6). Nucleotide diversity and other characteristics are summarised in supplemental Table 3. The patterns of ST distribution across geographical areas will be describe after (Fig. 3 and S7-S12).

**Figure 1.**
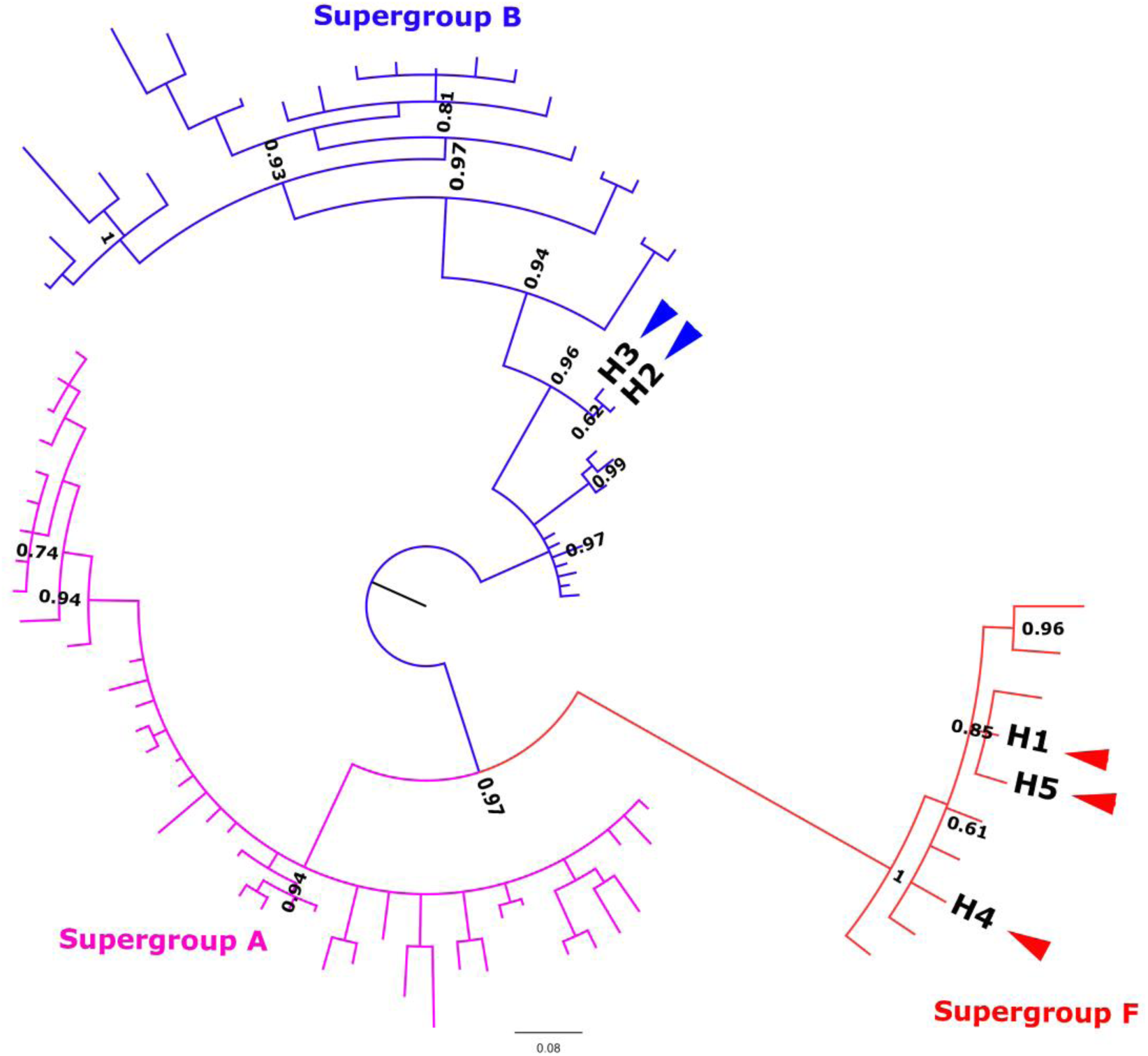
(Online colour figure) Summary unrooted phylogenetic tree of *fbpA alleles in several insects, including Chorthippus parallelus*, obtained by Bayesian inference. Alleles described in *C. parallelus* are named H1 to H5 *(marked as coloured arrows)*. Posterior probabilities are shown at the nodes. Other MLST genes are also analysed (Supplemental Fig. S2-S6). Sequence accession numbers are presented in Tables S13-S18.

**Figure 2.**
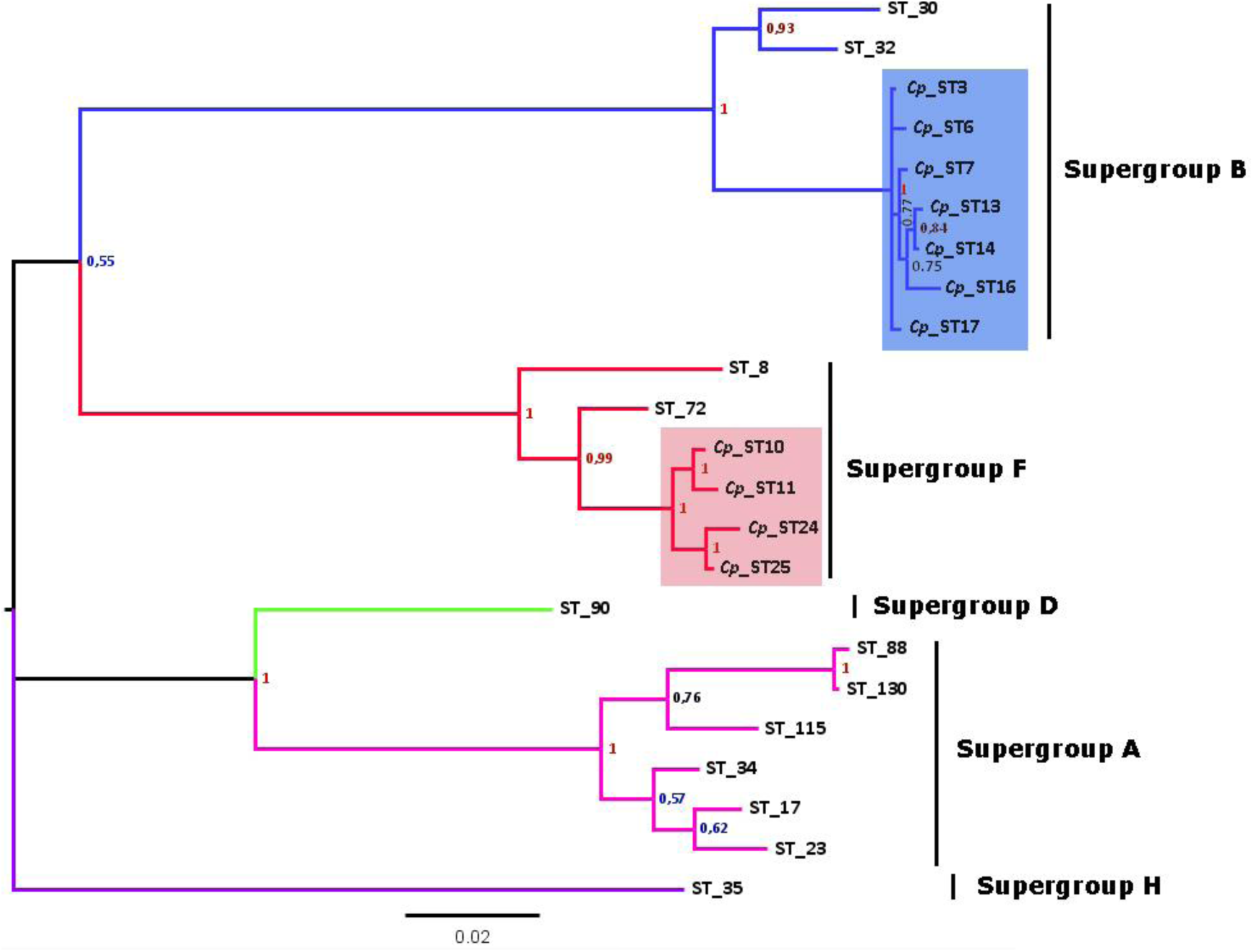
(Online colour figure) Phylogenetic tree of *Wolbachia* STs detected in *C. parallelus* (marked as Cp, coloured squares) excluding recombinants (see Fig. 4) obtained by Bayesian inference. The alleles described in this grasshopper bear the prefix Cp_ST. All other STs, named according to the official nomenclature, are available in the MLST database http://www.mlst.net/. Posterior probabilities are shown at the nodes.

**Figure 3.**
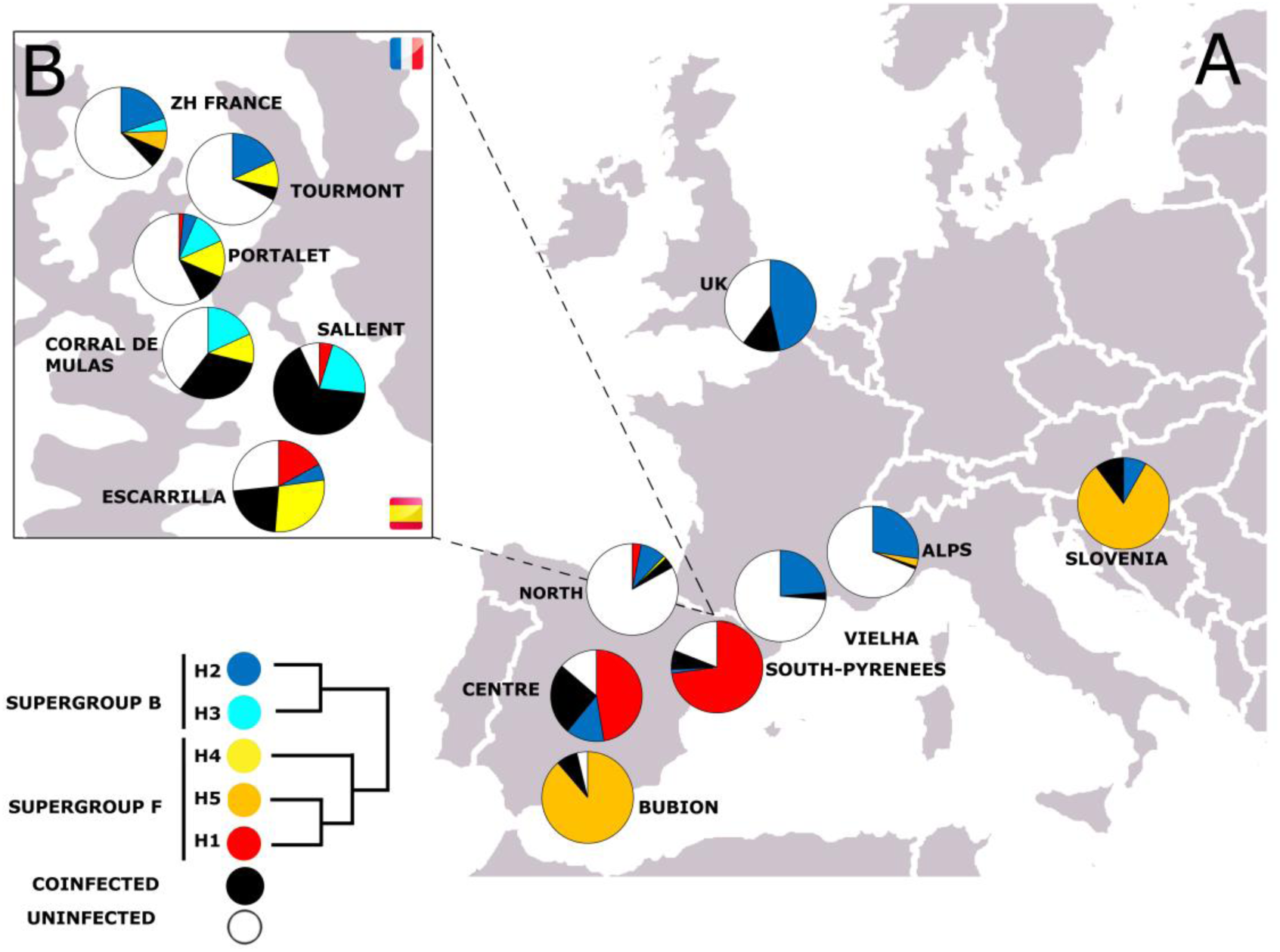
(Online colour figure) A) Geographical distribution of *fbpA* alleles in the *C. parallelus* populations analysed. Pyrenean hybrid zone (Tena’s valley, Huesca, Spain) is zoomed in B. See Table 1 for details.

**Table 3.**
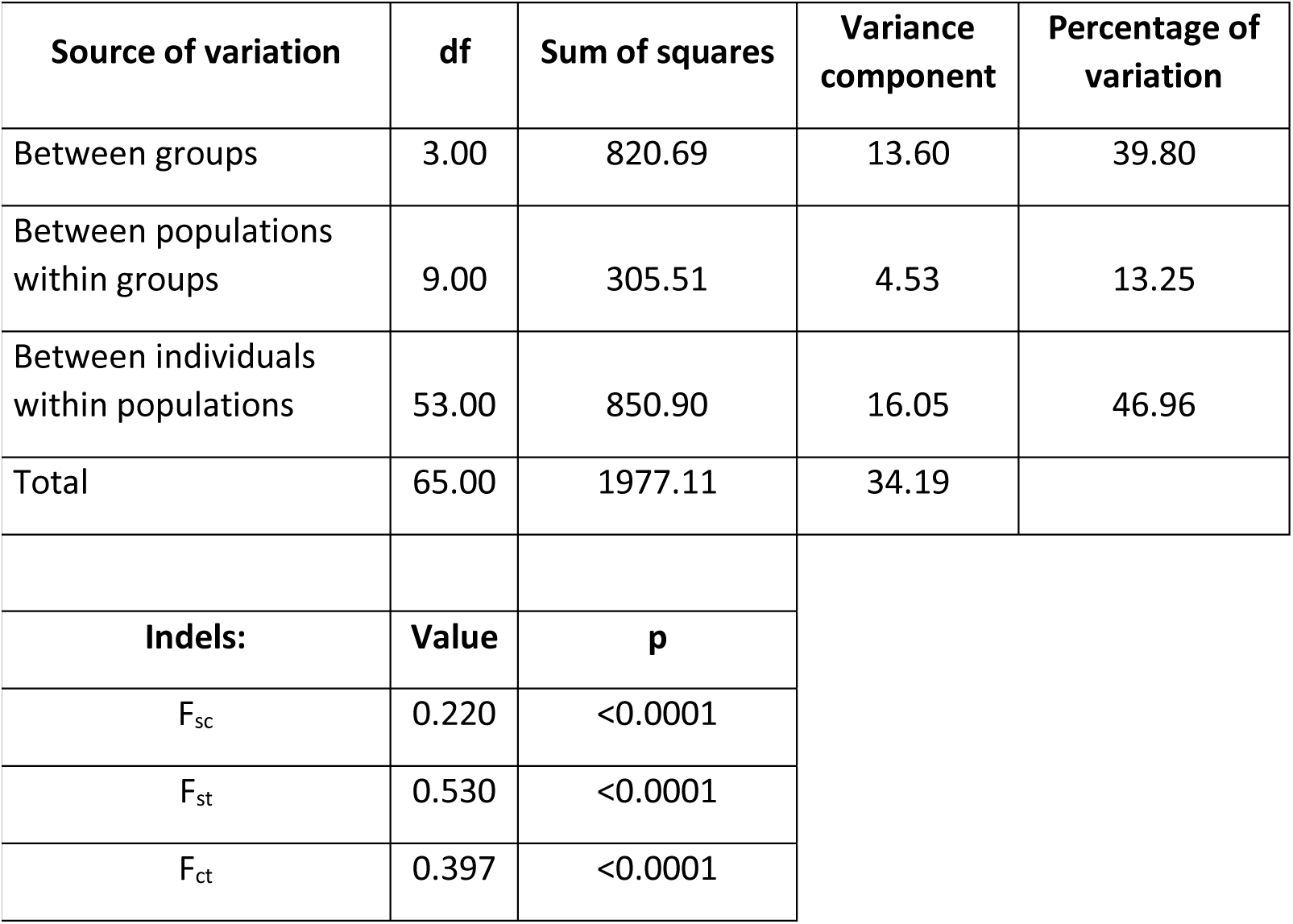
Analysis of molecular variance (AMOVA) from five MLST genes for the F supergroup of *Wolbachia* infecting different populations of *C. parallelus*.

| Source of variation | df | Sum of squares | Variance component | Percentage of variation |
| --- | --- | --- | --- | --- |
| Between groups | 3.00 | 820.69 | 13.60 | 39.80 |
| Between populations within groups | 9.00 | 305.51 | 4.53 | 13.25 |
| Between individuals within populations | 53.00 | 850.90 | 16.05 | 46.96 |
| Total | 65.00 | 1977.11 | 34.19 |  |
| **Indels:** | **Value** | **p** |  |  |
| F<sub>sc</sub> | 0.220 | <0.0001 |  |  |
| F<sub>st</sub> | 0.530 | <0.0001 |  |  |
| F<sub>ct</sub> | 0.397 | <0.0001 |  |  |

#### 1.2 Bacterial recombination

Recombination was detected in 19 STs or haplotypes. Recombinant *Wolbachia* strains were detected by two methods. Firstly, RDP4 analysis detected recombinant strains under several tests (marked “R” in Fig. 4, in contrast with parental strains, which are indicated as “F” or “B”, depending on the supergroup). In addition, the appearance of alleles of the B supergroup in isolates of the F supergroup (based on most of the genetic markers), and *vice versa*, also indicates recombination events. Our analysis revealed that both supergroups have exchanged parts of their genomes in some populations of *C. parallelus*, such as those of Portalet or Tourmont, in the centre of the hybrid zone, while recombination has not been detected in the grasshopper’s pure populations within or outside the hybrid zone. Some recombinants have also been detected in the north of Spain, in populations of this grasshopper characterised as hybrid on the basis of chromosomal markers (Bella *et al*. 2007).

**Figure 4.**
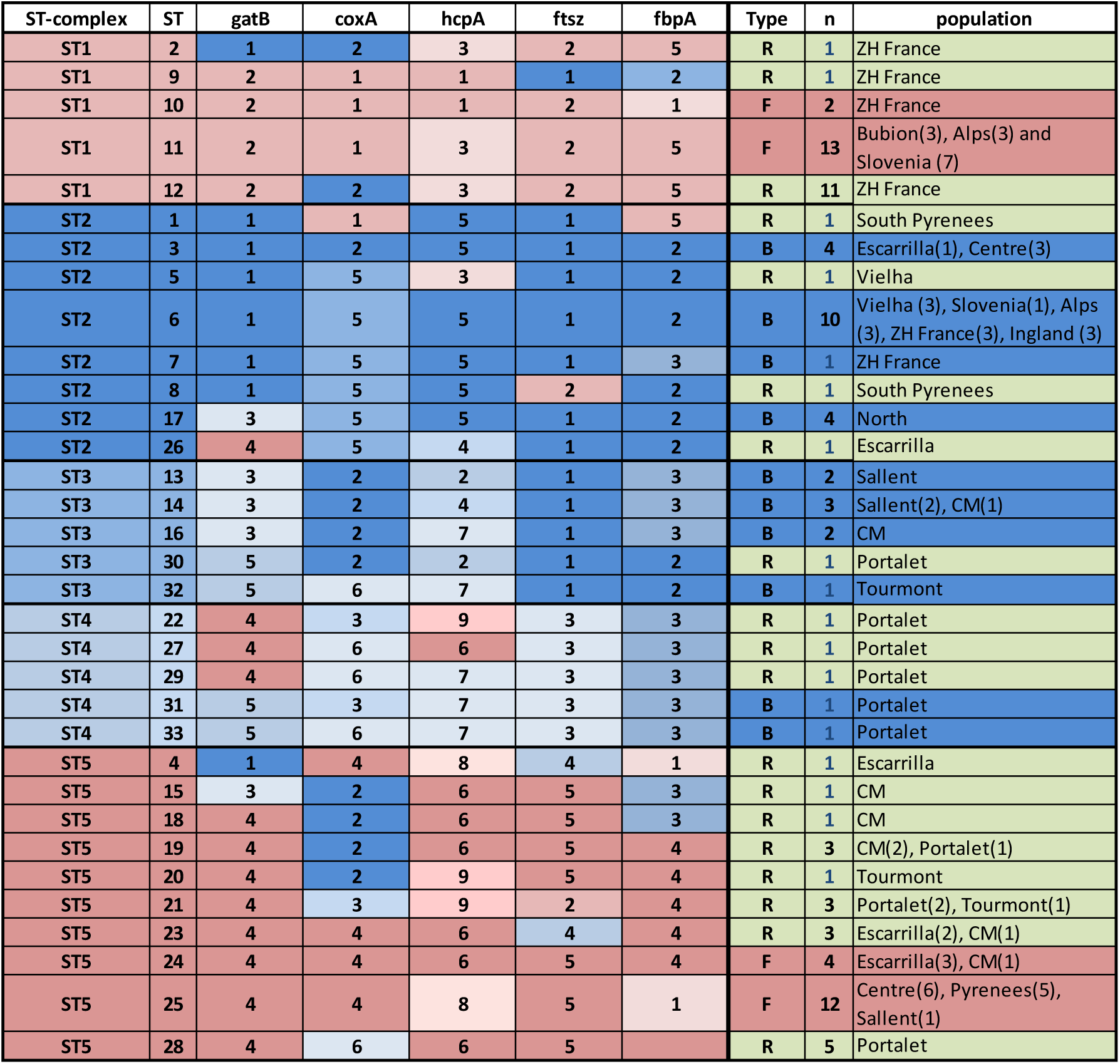
(Online colour figure) *Wolbachia* ST-complexes and allelic profiles described in *C. parallelus*. Note the classification in three groups: those assigned to supergroups F and B strains (“F” and “B”, respectively) and those in which possible recombination events between these supergroups were observed (“R”). Alleles belonging to F supergroup (see Fig. 1 and S2-S6) are marked with different red tones, while alleles belonging to the B supergroup are marked with blue tones. STs detected in only one individual (blue) should be interpreted with caution, even if the alleles appear in more than one sample. The name of population and number of individual (parenthesis) detected in each population are also indicated.

#### 1.3 Wolbachia phylogeny

After discarding recombinants STs to avoid artefacts, the phylogenetic tree from concatenated sequences allows distinguish 10 different strains of *Wolbachia* belonging to B supergroup and 4 different strains to F supergroup (Fig. 2).

In general, *Wolbachia* strains in *C. parallelus* are highly related. On the one hand, F strains are highly related between them. Other F strains, like *Wolbachia* infecting *Opistophthalmus granifrons (Scorpionida)* or *Cimex lectularius (Hemiptera)* are more distant. Their host is not related with *Chorthippus* (ecologically or phylogenetically). B strains are also related between them, but also with the *Wolbachia* strains infecting other Orthopteran, like *Teleogryllus taiwanemma*, and the recently detected *Wolbachia* strain infecting *Podisma pedestris*. This latest species shares habitat with *C. parallelus* (data not shown). Also, B strains (based on *16S rRNA* gene amplification) have been recently detected in other species, including *Ruspolia nitidula, Chorthippus vagans* and *Euhorthippus chopardi*, captured in the same populations that *Chorthippus parallelus* (Martinez-Rodriguez 2013).

Phylogenetic analyses of individual genes were also carried out. Alleles were correctly characterised as belonging to the F or B supergroups (Fig. 1 and Figs. S2-S6).

### 2 Biogeographical distribution of the Wolbachia strains

#### 2.1. *Individual* loci *analyses*

Different geographic isolates showed a high level of genetic variation within *Wolbachia* strains: Individual analysis of the 5 loci of MLST and the *wsp* gene allows us to determine a clear geographical pattern. In general, we detected alleles belonging to the B supergroup in *C. p. parallelus* and *C. p. erythropus* populations, indistinctly. By contrast, we can detect some alleles, belonging to F supergroup, specifically in some populations of *C. p. erythropus* or *C. p. parallelus* (see Fig. 3). In addition, we noted the presence of new alleles exclusive to *Wolbachia* infecting the hybrid grasshopper populations (Fig. 1, 3 and S2-S12). For instance and regarding gene *fbpA*, three alleles were identified belonging to supergroup F, and two alleles were assigned to supergroup B (Fig. 1). In the first case, allele 5 has been described in European populations of *C. parallelus* and Bubion in southern Spain and in the populations of the Cantabrian region (hybrid populations, *sensu* Bella *et al*. 2007). It has also been identified in the pure population of Gabas, on the French side of the hybrid zone (ZH France, Fig. 3). Allele 1, which also belongs to supergroup F, was detected in pure populations from the centre of the Iberian Peninsula and in the South Pyrenees populations of Escarrilla, Sallent and Portalet (hybrid zone). In the case of supergroup B, allele 2 was found in most of the populations. However, in the hybrid populations of Sallent, Corral de Mulas and Portalet (hybrid zone) we also detected alleles 3 and 4. Allele 4 has also been detected in Cantabrian hybrid populations. Similar patterns have been observed for the rest of the analyzed genes.

##### 2.1.1 *Wolbachia* ST- complexes

According with the MLST system implemented in START2 (Jolley *et al*. 2001), we classify the different haplotypes or ST in five ST-complexes (each one defined as a group of STs sharing a minimum of three alleles) (Fig. 4). The first ST-complex included several STs, some of them belonging to the F supergroup or recombinants highly related to this supergroup (see before, *Wolbachia* phylogeny), sampled in several non-Iberian populations from the rest of Europe and in some samples from the Basque Country in the north of Spain. The second ST complex, included isolates belonging to the B supergroup (and some recombinants, highly related to the B supergroup) widely distributed in *C. parallelus* populations, including both subespecies, but in different proportions. ST3 and ST4 complexes also include some recombinants and B strains. Finally, ST5 complex belongs to F supergroup but it shares some alleles with those of the B supergroup. They have been detected in *C. p. erythropus*, including some populations of the centre of Spain, and the Spanish region of the Pyrenees. The geographical distribution of each ST is showed in Fig. 4.

The ClonalFrame-based analysis infers *Wolbachia* microevolution using the multilocus sequence data and considers recombination. The genealogies confirmed the genetic subdivisions in the strains of the F supergroup (Fig. 5), while B strains were grouped in the same clade. The genealogies also detected the recombinant strains that mostly appear in the grasshopper hybrid zone. The clades also support an association between genetic and geographic data.

**Figure 5.**
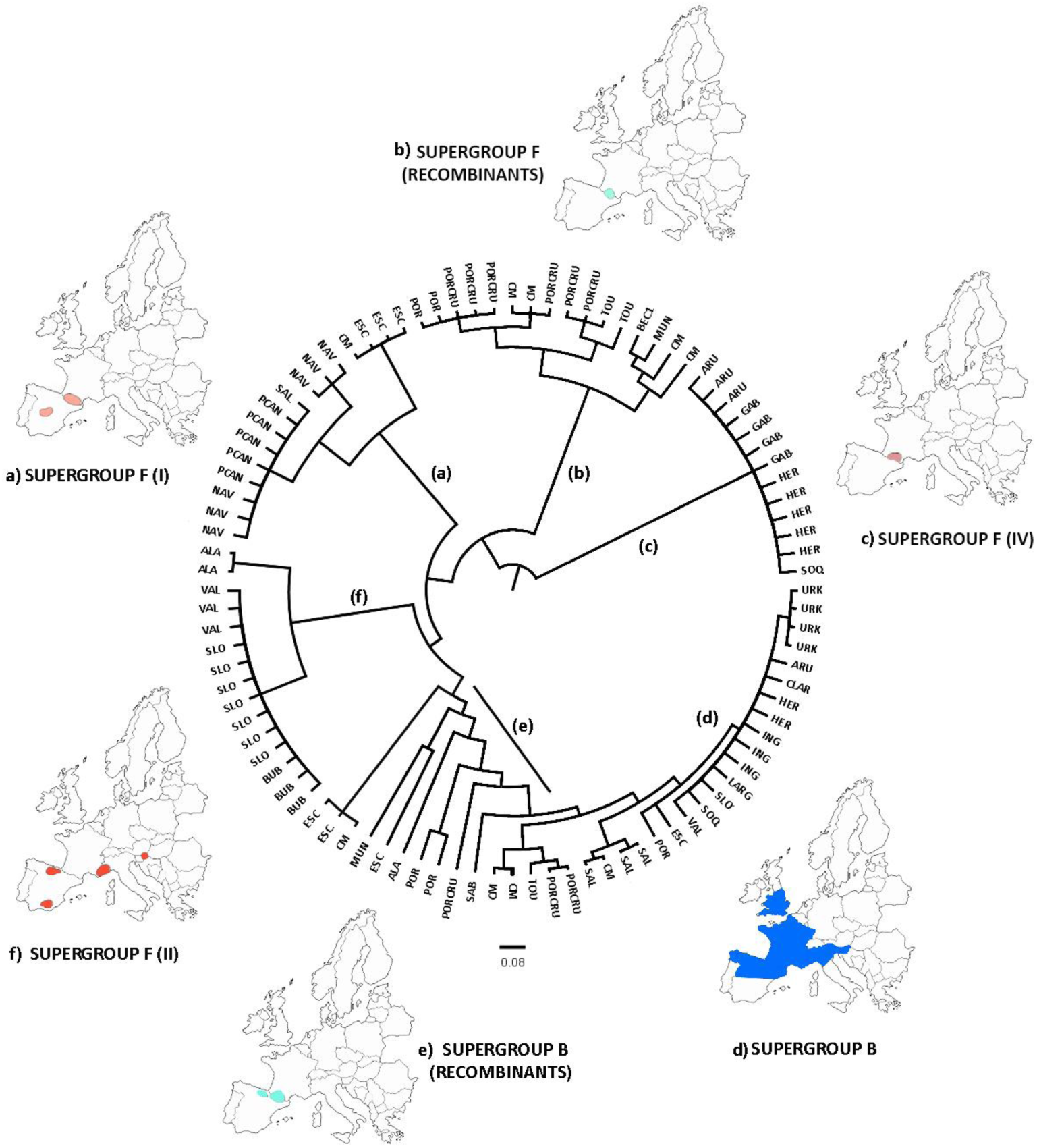
ClonalFrame genealogy (Online colour figure). Maps indicate the approximate location of the samples assigned to the major clades, classified with respect to their corresponding F or B supergroup. The analysis distinguished three major clades of supergroup F (a, c and f), one clade belonging to supergroup B (d), and several recombinant strains (b and e). This was consistent with our previous analyses. Acronyms are listed in Table 1.

The lower differentiation of isolates within geographic populations and the higher differentiation of those between geographic populations suggest isolation-by-distance between the bacterial F strains infecting the two grasshopper subspecies (with the exception of Bubion; see Discussion). The AMOVA indicated a geographic division of the F supergroup: (i) Central Iberian Peninsula and South Pyrenees populations, (ii) Pyrenean hybrid zone populations, (iii) French side of the hybrid zone, and (iv) Non-Iberian populations from the rest of Europe and Bubion (in Spain) (Table 3).

These results were also supported by the locus-by-locus AMOVA (except for *hcpA*) (Table S4) and the exact test of population differentiation (Rousset *et al*. 1992) (Table S5).

In addition, the Mantel tests confirmed that the genetic and geographic distances were correlated (rY1: 0.338, p=0.001). This correlation was stronger when the Bubion data were excluded (rY1: 0.483, p=0.003). This particular geographical distribution could be related to the biogeographical distribution of this grasshopper during the last glaciation and allows us to infer the origin of *Wolbachia* infection in *C. parallelus* and its role in establishing the hybrid zone.

### 3 Estimation of Wolbachia divergence dates

Previous studies suggest a synonymous divergence rate of about 0.90% per million years (MY) for bacteria. However, this bacterial molecular clock should be interpreted with caution since divergence rates may differ between bacteria species (Ochman *et al*. 1999; Ochman & Wilson 1987; Raychoudhury *et al*. 2009). Based on this estimate, the divergence between the F strains detected in the centre of Spain (Cp_ST-25) and Slovenia (Cp_ST-11) is about 3,400,000 years. On the other hand, the divergence between the F strains detected in the centre of Spain (Cp_ST-25) and the hybrid zone (Cp_ST-24) is about 1,400,000 years. B strains detected in the centre of Spain (Cp_ST-3) and the widely distributed Cp_ST-6 diverged about 250,000 years ago. The dates of divergence of strains based on the different markers are illustrated in supplementary Tables S6-S12. Substitution rate could represent a lower boundary for the mutation rate within strains (Emerson 2007). Thus, other estimation of intraspecific mutation rate (in terms of *D. melanogaster* generations (6.87E-10 per position per insect generation in the 3rd position, see Richardson *et al*. 2012) have been used. Divergence between strains could be higher (x 10) if we consider this estimation of *Wolbachia* evolutionary rates.

## Discussion

### Modes of acquisition of Wolbachia. Codivergence *vs.* horizontal transmission

Three hypotheses about the origin of *Wolbachia* in *Chorthipus parallelus* are discussed: Firstly, an ancient co-divergence between *Wolbachia* and this orthopteroid. Secondly, the acquisition of *Wolbachia* before the subspecies divergence, and the recent co-divergence of *Wolbachia* and the host. And thirdly, the recent acquisition of *Wolbachia* by horizontal transmission.

*Wolbachia* codivergence with their host is extremely rare in the literature compared with horizontal acquisition between species (Raychoudhury *et al*. 2009). To distinguish between co-divergence and horizontal transmission events, a good knowledge of the recent evolutionary history of the host is required as it happens with the *C. parallelus* system which becomes a good model to study *Wolbachia* expansion. Our *Wolbachia* phylogenetic and phylogeographic data can also be interpreted in the context of its host evolution so serving to infer the *Wolbachia* transmission and evolution in this particular grasshopper and its influence in the hybrid zone.

Not discarding other mechanisms that could also be involved (some paternal transmission, infection loss, drive, etc.), our data point out two possible mechanisms to explain current *Wolbachia* infection in both *Chorthippus parallelus* subspecies: the codivergence of *Wolbachia* F strains during recent speciation of both subspecies followed by “modern” horizontal transmission of B strains from other organisms.

#### a) Phylogenetic relationships *&* common biogeography between host and bacteria

Phylogenetic data support that bacterial F strains infecting *Chorthippus* are extremely similar among them, and that they are highly related each other than with any other outside this grasshopper’s taxa (see Fig. 1, 2 and supplemental figures S2-S6). These data support the recent co-divergence between host and bacteria. Furthermore, the geographical distribution of two main F bacterial lineages is largely congruent with the biogeography of *C. parallelus* (Lunt et al; 1998). Cp25 and Cp24 linages infect *C. p. erythropus*, while Cp11 infects mainly *C. p. parallelus* (except the Iberian southern population of *C. p. erythropus* of Bubion). Hybrid grasshoppers are infected by variants of both lineages. Strains geographical distribution supports the co-divergence between the two subspecies and the two main F strains infecting *C. parallelus*.

However, the co-divergence between *Wolbachia* and their host should be “recent”. F supergroup (based on *16S rRNA* and *Ftsz* genes) has been detected in the bush cricket species *Orocharis saltator* and *Hapithus agitator* (Gryllidae: Eneopterinae) but no in other Acrididae (both families have diverged 300 Ma ago; see Song *et al*. 2015). In addition, both F *Wolbachia* infecting *Chorthippus* are closer to strains infecting other insect orders than to this Gryllidae one. This suggests that the F strain of *Wolbachia* was acquired by horizontal transmission before the divergence between subspecies, followed by co-divergence between each host and bacteria in their corresponding glacial refugia and during postglacial expansion.

By contrast, B supergroup strains infect homogeneously both subspecies, without a biogeographical pattern. The variability within B supergroup is restricted to the hybrid zone, in which new variants and alleles, highly related, appear. All data suggest a recent and quick horizontal transmission of B strain to this host.

#### b) Divergence time estimation

Our current data serve to estimate the divergence time of *Wolbachia* according with a general bacterial molecular clock (Ochman *et al*. 1999; Ochman & Wilson 1987; Raychoudhury *et al*. 2009). This supports that the divergent time of *Wolbachia* F strains is higher (3.4-1.4 Myr) than *C. parallelus* subspecies divergence time (500.000 years according with mtDNA data, Hewitt, 1996). However, this estimation could represent a lower boundary for the mutation rate within species (Emerson 2007). Furthermore, we have also estimated this time of *Wolbachia* divergence higher (x10) according with some specific *Wolbachia* mutation rates noticed in *Drosophila* (Richardson *et al*., 2012). However, several factors can lead to inappropriate estimation of divergence dates. For instance, this estimation in *Drosophila* could be inappropriate for *Chorthippus*, in which each host generation takes one year, which modifies the dynamic of *Wolbachia* transmission, not discarding possible bottlenecks of bacterial population, selection pressures, etc. Due to that, we think that the divergence times are compatible, even when they are not coincident, and support an ancient acquisition of F *Wolbachia*, followed by its co-divergence with their *Chorthippus* hosts.

By contrast, B *Wolbachia* divergence times are lower, and suggest a more “recent” acquisition by *C. parallelus* by horizontal transmission. The existence of an extremely close B strain of *Wolbachia* in a number of orthopteran species that share the same habitats also support this hypothesis (Martinez-Rodriguez, 2013):

#### c) Horizontal transmission from other taxa

We have detected extremely closely related B strains in other orthopteroids like *Podisma pedestris, Chorthippus vagans* and *EuChorthippus chopardi* (Acrididae) but also *Ruspolia nitidula* <u>(Tettigoniidae)</u> that share habitat with *Chorthippus* (data no shown, Martinez-Rodriguez, 2013). Most genera belonging to family Acrididae diverged 50 Ma ago, and both families (Acrididae and Tettigoniidae) did it 250 Ma ago (Song *et al*. 2015). The incongruence between *Wolbachia* and host divergence times supports that the B supergroup could have been “recently” acquired as a result of rapid expansion of the infection from other taxa (horizontal transmission). In this context, some species of parasitoids could be a vector for intra- or inter-specific infection transmission (unpublished data, Martinez-Rodriguez, 2013).

By contrast, a recent horizontal transmission of F strain is unlikely. There is no evidence of closely F *Wolbachia* strains currently infecting other Orthoptera. However, F strain (usual mutualist of nematodes, but also present in arthropods) could infect another insect in the past and explain the horizontal transmission of *Wolbachia* to an ancestral *C. parallelus* before subspecies divergence. Even if we cannot totally discard a recent horizontal transmission, we consider that the hypothesis of 2 independent “recent” acquisitions of 2 related F strains to this 2 geographically distant subspecies of *Chorthippus* is unlikely. In our opinion, the hypothesis of an “ancient” acquisition (>4 Myr) and consecutive co-divergence of *Wolbachia* F strains is more likely.

This hypothesis is also supported by the detection of an insertion of *Wolbachia* that coincides in homologous chromosomes of both Cpe and Cpp, while other inserts are subspecies-specific (Funkhouser-Jones *et al*. 2015, Toribio-Fernandez *et al*., *in. prep*.). This recent finding supports that an ancestral *Chorthippus* sp. was already infected by *Wolbachia*, before divergence of two subspecies.

### Diversification of Wolbachia inside the hybrid zone

The ST distribution suggests that there is a particular pattern of *Wolbachia* infection within the Pyrenean grasshopper hybrid zone and suggests that the particular interaction between host “hybrid genomes” and bacterial infection could happen. B and F supergroups are in contact in several populations of *Chorthippus parallelus*, to the point of coexisting in the same individuals (coinfection, see below). However, we only have detected these new, recombinant strains in hybrid populations (inside the Pyrenees hybrid zone but also in a hybrid population in northern Spain).

High recombination between *Wolbachia* strains has been reported several times (Foster *et al*. 2011; Jiggins 2002; Jiggins *et al*. 2001; Verne *et al*. 2007; Werren & Bartos 2001). In fact, recombination levels in *Wolbachia* seem to be higher than, for example, in *Neisseria meningitidis*, which is considered a bacterium with a great capacity for recombination (Jolley et al. 2005). However, different recombination rates between strains have been detected (Klasson *et al*. 2009). The recombination process serves the strains to vary and adapt rapidly, which is important for their interaction with the host. For instance, data show that mutualist strains, adapted to a particular host, have limited levels of recombination compared with other strains than potentially should adapt to a new host (Jiggins 2002; Werren & Bartos 2001). Actually, the preservation of a high number of genes of recombination guarantees genomic flexibility during recurrent host change (Darby *et al*., 2007; Hurst *et al*., 2002).

This hypothesis is pertinent to the case of *Wolbachia* strains that infect hybrid *C. parallelus*. The contact of bacteria with in a new host (the hybrid grasshoppers) could have resulted in a high bacterial recombination rate in order to adapt to this new host. It might explain why our analyses detect recombinant strains infecting grasshoppers just in hybrid populations, although the F and B supergroups are in contact in many other *C. parallelus* populations (Bella *et al*. 2010; Zabal-Aguirre *et al*. 2010; Zabal-Aguirre *et al*. 2014).

In addition, we also detected, specifically in the grasshopper hybrid zone, new bacterial alleles belonging to these recombinant strains, which have diverged from closely related B and F alleles found in other isolates. This suggests that sequences diverged rapidly after recombination. Possible explanations include that *Wolbachia* strains infecting grasshopper hybrids diverged separately of other strains (due to the isolation between hybrids and pure populations) or perhaps these strains support other evolutionary pressures (for instance, adaptation to other a new hybrid host or to the evolutionary processes involved in the hybrid zone). More studies will be needed in order to clarify this.

### The origin and expansion of Wolbachia infection in C. parallelus and its effects on the dynamic of the hybrid zone

We consider that recent codivergence is the best explanation for F *Wolbachia* strains appearance in both subespecies of *C. parallelus*, while B infection is better explained by modern horizontal transmission. According with that, we propose a possible scenario to explain the *C. parallelus Wolbachia*’s acquisition (Fig. 6).

**Figure 6.**
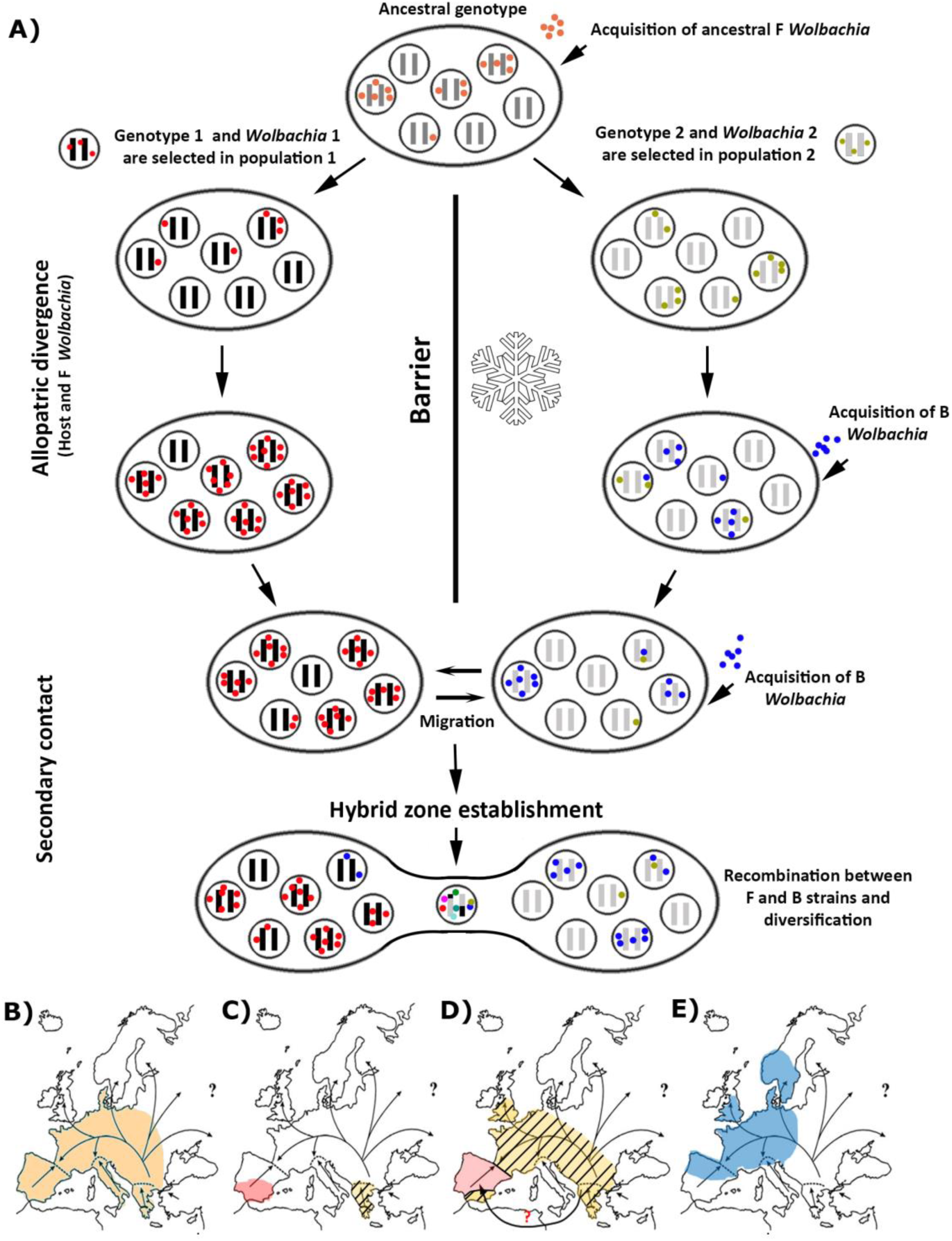
(Online colour figure). A) Proposed hypothesis for the origin of *Wolbachia* infection in *C. parallelus*. Each ellipse represents a population. Inner circles represent individuals. Black and grey bars indicate the host genome, while the coloured dots show the bacterial type infecting the individual. The hybrid zone would be established simultaneously with the appearance of recombinant genomes in the host, and a high bacterial diversity, induced by recombination. B) Spatial representation of the population expansion of infection: the arrows indicate the population expansion of *C. parallelus* (modified from Hewitt 2001), after the retreat of the glacial ice. Before the last glaciation the infection of *Wolbachia* by the F supergroup was homogeneous. C) During the last glaciation, *C. parallelus* and F *Wolbachia* diverged in allopatry. D) After the ice disappeared, the pattern of expansion of the F infection coincided with that of the migration of its host, E) Recently, B infection has been transmitted horizontally in different European populations.

The study of *Wolbachia* infecting *C. parallelus* divergence should be considered in the context of the last quaternary ice age in Europe and its consequences for *C. parallelus* distribution: During this glaciation the grasshopper subspecies diverged as a result of their geographic isolation in allopatry (Hewitt 1993, 1996, 1999, 2001, 2011; Serrano *et al*. 1996). After the retreat of the ice, grasshopper populations from the Iberian Peninsula colonised the Pyrenees, meeting *C. p. parallelus* coming from the Balkans, as suggested by Lunt *et al*. (1998).

The current data about *Wolbachia* infection suggest that an ancient F strain of *Wolbachia* and an ancestral host could codiverge during this period before meeting when the hybrid zone formation. In addition, a new B infection could have been acquired more recently, and expanded in the pure and hybrid populations afterwards. *Wolbachia* spread by horizontal transmission can be very effective, as previously suggested (Turelli & Hoffmann, 1991). In addition, the lack of B infection in some populations, like Bubion in Southern Spain, also points out a recent spread of infection from continental Europe (where it is massive): the isolation of these individuals and their geographical location has not permitted their infection yet. Loss of an ancestral B infection in this population seems less plausible to us, given the strain’s aforementioned homogeneity and abundance.

Finally, after the hybrid zone formation, new strains would have arisen in the hybrid zone by recombination. We are reminded that the appearance of the F strains in the grasshopper populations of central and southern Spain could be explained by an alternative route of colonization (from the East or the South), not discarding either other less parsimonious hypotheses.

### *Wolbachia* effects in the hybrid zone: New perspectives

Our data suggest that *Wolbachia* already infected *C. parallelus* during the hybrid zone formation. Due to that, *Wolbachia*’s role in the hybrid zone dynamic deserves some discussion: We propose that genetic incompatibilities between the grasshopper subspecies accumulated during the divergence, together with the unidirectional and bidirectional CI that *Wolbachia* induces in the hybrid zone (Zabal-Aguirre *et al*., 2014) thereby influencing the formation of the current grasshopper hybrid zone. More data are required to quantify the importance of CI in hybrid formation.

In the other hand at least two F strains of *Wolbachia* infect differently *C. parallelus* subspecies. New experiments should be carried out in order to verify if further CI exists, induced within those strains belonging to F supergroup. It is possible that these new bacterial lineages or STs, the result of processes of recombination between strains from supergroups F and B, and their subsequent diversification by point mutations and adaptation, would limit the incompatibility between grasshopper individuals infected by different supergroups in hybrid populations, favouring the appearance of a new, mixed bacterial and host genetic background in this area, in contrast to the pure populations on either side of the Pyrenees. This new scenario should be tested to know the current and actual role of *Wolbachia* in the *C. parallelus* hybrid zone and for a better study of this model of incipient speciation.

## Acknowledgments

We are grateful to the authorities that gave us permission to collect the specimens analysed here: Parc National des Pyrenèes, Parque Nacional de Sierra Nevada, Comunidad Autónoma de Madrid, Comunidad Autónoma de Aragón, Comunidad Autónoma de Castilla y León, Parc National du Mercantour Alpes-Maritimes. Dr PL Mason (University of Glasgow, UK) helped us with the manuscript and Dr JR Dagley and Dr R Kostanjšek kindly provided the individuals from the Epping Forest (UK) and Mokronog (Slovenia), respectively. We also thank other members of our HYZO group in the Universidad Autónoma de Madrid, who helped with insect collection and handling, or contributed to discussions about our study. This work has been supported by the Spanish MINECO I+D+i grants CGL2009-08380/BOS and CGL2012-35007 and the collaboration of Chromacell.

## Data accessibility

DNA sequences: GenBank database under accession numbers KM078849-KM078883

## Author contribution

Martinez-Rodriguez, P designed research, performed research, analyzed data and wrote the paper. Arroyo-Yebras, F performed research. Bella, JL designed research, analyzed data and wrote the paper. All of them, with the contribution in some cases of other members of the group and the collaborators cited in the acknowledgments section, collected the grasshoppers.

## Supplemental tables

**Table S1.** *Accession numbers*.

| gen | allele | Genebank code |
| --- | --- | --- |
| CoxA | H1 | KM078848 |
|  | H2 | KM078849 |
|  | H3 | KM078850 |
|  | H4 | KM078851 |
|  | H5 | KM078852 |
|  | H6 | KM078853 |
| fbpA | H1 | KM078854 |
|  | H2 | KM078855 |
|  | H3 | KM078856 |
|  | H4 | KM078857 |
|  | H5 | KM078858 |
| ftsZ | H1 | KM078859 |
|  | H2 | KM078860 |
|  | H3 | KM078861 |
|  | H4 | KM078862 |
|  | H5 | KM078863 |
| gatB | H1 | KM078864 |
|  | H2 | KM078865 |
|  | H3 | KM078866 |
|  | H4 | KM078867 |
|  | H5 | KM078868 |
| hcpA | H1 | KM078869 |
|  | H2 | KM078870 |
|  | H3 | KM078871 |
|  | H4 | KM078872 |
|  | H5 | KM078873 |
|  | H6 | KM078874 |
|  | H7 | KM078875 |
|  | H8 | KM078876 |
|  | H9 | KM078877 |
|  | H10 | KM078878 |
| wsp | H1 | KM078879 |
|  | H2 | KM078880 |
|  | H3 | KM078881 |
|  | H4 | KM078882 |
|  | H5 | KM078883 |

**Table S2.** *Wolbachia* infection frequencies in the analysed populations:

| NAME | n |  |  |  |  | % |  |  |  |  |
| --- | --- | --- | --- | --- | --- | --- | --- | --- | --- | --- |
|  | Ø | F | B | FB | total | Ø | F | B | FB | TOTAL |
| ESC2008 | 19 | 7 | 3 | 9 | 38 | 0,50 | 0,18 | 0,08 | 0,24 | 1,00 |
| ESC2009 | 42 | 34 | 5 | 7 | 88 | 0,48 | 0,39 | 0,06 | 0,08 | 1,00 |
| SAL2008 | 3 | 3 | 12 | 30 | 48 | 0,06 | 0,06 | 0,25 | 0,63 | 1,00 |
| SAL2009 | 9 | 5 | 16 | 21 | 51 | 0,18 | 0,10 | 0,31 | 0,41 | 1,00 |
| CM2008 | 41 | 9 | 0 | 4 | 54 | 0,76 | 0,17 | 0,00 | 0,07 | 1,00 |
| CM2009 | 25 | 7 | 10 | 17 | 59 | 0,42 | 0,12 | 0,17 | 0,29 | 1,00 |
| POR2008 | 36 | 14 | 2 | 9 | 61 | 0,59 | 0,23 | 0,03 | 0,15 | 1,00 |
| POR2009 | 69 | 12 | 10 | 0 | 91 | 0,76 | 0,13 | 0,11 | 0,00 | 1,00 |
| C_TOU2008 | 45 | 5 | 9 | 1 | 60 | 0,75 | 0,08 | 0,15 | 0,02 | 1,00 |
| C_TOU2009 | 46 | 4 | 10 | 3 | 63 | 0,73 | 0,06 | 0,16 | 0,05 | 1,00 |
| SOQ2008 | 20 | 2 | 7 | 3 | 32 | 0,63 | 0,06 | 0,22 | 0,09 | 1,00 |
| SOQ2009 | 27 | 3 | 18 | 11 | 59 | 0,46 | 0,05 | 0,31 | 0,19 | 1,00 |
| LHER2008 | 24 | 4 | 15 | 5 | 48 | 0,50 | 0,08 | 0,31 | 0,10 | 1,00 |
| LHER2009 | 23 | 6 | 16 | 10 | 55 | 0,42 | 0,11 | 0,29 | 0,18 | 1,00 |
| GAB2008 | 30 | 6 | 6 | 3 | 45 | 0,67 | 0,13 | 0,13 | 0,07 | 1,00 |
| GAB2009 | 36 | 9 | 10 | 8 | 63 | 0,57 | 0,14 | 0,16 | 0,13 | 1,00 |
| ARU2008 | 20 | 1 | 13 | 0 | 34 | 0,59 | 0,03 | 0,38 | 0,00 | 1,00 |
| ARU2009 | 71 | 2 | 9 | 8 | 90 | 0,79 | 0,02 | 0,10 | 0,09 | 1,00 |
| NAV2007 | 22 | 36 | 17 | 33 | 108 | 0,20 | 0,33 | 0,16 | 0,31 | 1,00 |
| NAV2008 | 0 | 8 | 4 | 10 | 22 | 0,00 | 0,36 | 0,18 | 0,46 | 1,00 |
| PCANTO2008 | 21 | 22 | 0 | 6 | 49 | 0,43 | 0,45 | 0,00 | 0,12 | 1,00 |
| MUNA2008 | 21 | 5 | 19 | 7 | 52 | 0,40 | 0,10 | 0,37 | 0,14 | 1,00 |
| COLLARCHE2009 | 16 | 0 | 5 | 0 | 21 | 0,76 | 0,00 | 0,24 | 0,00 | 1,00 |
| VALDIERI I and II 2009 | 55 | 3 | 17 | 1 | 76 | 0,72 | 0,04 | 0,22 | 0,01 | 1,00 |
| LARGENTERA2009 | 2 | 0 | 6 | 0 | 8 | 0,25 | 0,00 | 0,75 | 0,00 | 1,00 |
| BUBION2004 | 2 | 47 | 0 | 4 | 53 | 0,04 | 0,89 | 0,00 | 0,08 | 1,00 |
| EPPING2002 | 6 | 0 | 7 | 2 | 15 | 0,40 | 0,00 | 0,47 | 0,13 | 1,00 |
| VIELHA2008* | 28 | 8 | 1 | 1 | 38 | 0,74 | 0,21 | 0,03 | 0,03 | 1,00 |
| URKAREGUI2009 | 35 | 7 | 6 | 1 | 49 | 0,71 | 0,14 | 0,12 | 0,02 | 1,00 |
| TXABARRI2009 | 9 | 0 | 0 | 1 | 10 | 0,90 | 0,00 | 0,00 | 0,10 | 1,00 |
| TARA2008 | 57 | 2 | 6 | 4 | 69 | 0,83 | 0,03 | 0,09 | 0,06 | 1,00 |
| TARA2009 | 73 | 1 | 0 | 3 | 77 | 0,95 | 0,01 | 0,00 | 0,04 | 1,00 |
| ALAV_1_2009 | 33 | 12 | 1 | 1 | 47 | 0,70 | 0,26 | 0,02 | 0,02 | 1,00 |
| SLOVENIA2009 | 0 | 40 | 4 | 5 | 49 | 0,00 | 0,82 | 0,08 | 0,10 | 1,00 |
| TOTAL | 966 | 324 | 264 | 228 | 1782 | 0,54 | 0,18 | 0,15 | 0,13 | 1,00 |

**Table S3.** Genetic diversity:

| Gene | n | P | S | Eta | K | Hap | Hd/VarHd | Pi | ThetaNuc | AvNumDif | ThetaG | TajimaD | FuLiD* | FuLiF* | G+C | R<br>(MAXCHI,<br>p<0.01) |
| --- | --- | --- | --- | --- | --- | --- | --- | --- | --- | --- | --- | --- | --- | --- | --- | --- |
| *coxa* | 111 | 402 | 41 | 42 | 16.25 | 6 | 0.8/0.0002 | 0.040 | 0.020 | 16.251 | 7.951 | 3.238** | 2.122** | 3.090** | 0.385 | No |
| *fbpa* | 117 | 429 | 61 | 62 | 11.62 | 5 | 0.8/0.0001 | 0.068 | 0.028 | 28.579 | 11.621 | 4.625*** | 2.307** | 3.943** | 0.394 | Yes: 3<br>(83) |
| *ftsZ* | 112 | 435 | 59 | 59 | 27.80 | 5 | 0.7/0.0002 | 0.064 | 0.026 | 27.801 | 11.151 | 4.740*** | 2.274** | 3.965** | 0.407 | Yes: 1<br>(34) |
| *gatB* | 114 | 370 | 39 | 39 | 17.30 | 5 | 0.7/0.0004 | 0.047 | 0.020 | 17.298 | 7.346 | 4.165*** | 2.092** | 3.517** | 0.370 | No |
| *hcpA* | 115 | 419 | 57 | 57 | 23.66 | 10 | 0.9/0.0002 | 0.056 | 0.026 | 23.662 | 10.719 | 3.815*** | 2.057** | 3.351** | 0.366 | No |

**Table S4.**
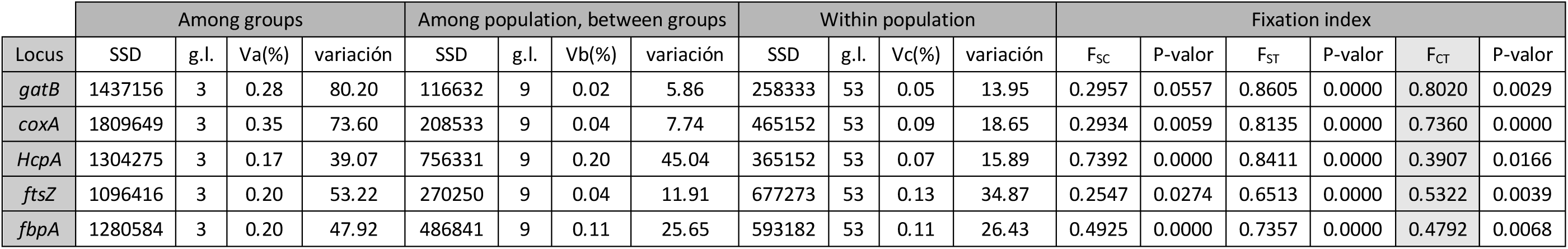
Locus by locus AMOVA implemented in ARLEQUIN.

**Table S5.** Exact test of population differentiation following the methodology of Rousset *et al*. (1992) implemented in ARLEQUIN. Gray = populations among which differentiation is observed.

|  | Pto. Cantó | Centre | Escarrilla | Sallent | Alps | North | Vielha | Bubió | Slovenia | Tourmon t | Portalet | C. Mulas | ZH Franc e |
| --- | --- | --- | --- | --- | --- | --- | --- | --- | --- | --- | --- | --- | --- |
| Pto. Cantó |  |  |  |  |  |  |  |  |  |  |  |  |  |
| Centr e | 0.00458 ± 0.0008 |  |  |  |  |  |  |  |  |  |  |  |  |
| Escar rilla | 0.10008 ± 0.0021 | 0.00216 ± 0.0005 |  |  |  |  |  |  |  |  |  |  |  |
| Sallen t | 0.24378 ± 0.0048 | 0.00015 ± 0.0001 | 0.10440 ± 0.0042 |  |  |  |  |  |  |  |  |  |  |
| Alps | 0.06187 ± 0.0015 | 0.00021 ± 0.0001 | 0.03494 ± 0.0017 | 0.39706 ± 0.0041 |  |  |  |  |  |  |  |  |  |
| North | 0.10136 ± 0.0015 | 0.00207 ± 0.0004 | ≈ 0.00000 | 0.10483 ± 0.0028 | 0.03525 ± 0.0011 |  |  |  |  |  |  |  |  |
| Vielh a | 0.01245 ± 0.0010 | 0.00002 ± 0.0000 | 0.01049 ± 0.0008 | 0.00210 ± 0.0006 | 0.00247 ± 0.0005 | 0.01230 ± 0.0006 |  |  |  |  |  |  |  |
| Bubió n | 0.05564 ± 0.0030 | ≈ 0.00000 | 0.02453 ± 0.0024 | 0.03982 ± 0.0011 | 0.00212 ± 0.0004 | 0.02717 ± 0.0008 | 0.00013 ± 0.0001 |  |  |  |  |  |  |
| Slove<br>nia | 0.50524 ±<br>0.0020 | 0.15809 ±<br>0.0027 | 0.25167 ±<br>0.0021 | 0.70584 ±<br>0.0059 | 0.27881 ±<br>0.0042 | 0.24826 ±<br>0.0023 | ≈ 0.00000 | 0.42498 ±<br>0.0123 |  |  |  |  |  |
| Tour<br>mont | 0.01702 ±<br>0.0008 | 0.00038 ±<br>0.0002 | 0.01731 ±<br>0.0006 | 0.01048 ±<br>0.0013 | 0.00451 ±<br>0.0004 | 0.01802 ±<br>0.0009 | ≈ 0.00000 | 0.00146 ±<br>0.0004 | ≈ 0.00000 |  |  |  |  |
| Portal<br>et | 0.00802 ±<br>0.0009 | ≈ 0.00000 | ≈ 0.00000 | 0.00049 ±<br>0.0003 | 0.00033 ±<br>0.0002 | ≈ 0.00000 | 0.00052 ±<br>0.0001 | 0.00046 ±<br>0.0003 | 0.12316 ±<br>0.0015 | 0.00135 ±<br>0.0003 |  |  |  |
| C.<br>Mula<br>s | 0.39703 ±<br>0.0045 | 0.03246 ±<br>0.0018 | 0.09899 ±<br>0.0013 | 0.57693 ±<br>0.0065 | 0.14184 ±<br>0.0023 | 0.10019 ±<br>0.0014 | 0.03558 ±<br>0.0014 | 0.37234 ±<br>0.0104 | 1.00000 ±<br>0.0000 | 0.04496 ±<br>0.0012 | 0.02705 ±<br>0.0015 |  |  |
| ZH<br>Franc<br>e | 0.49858 ±<br>0.0032 | 0.15600 ±<br>0.0035 | 0.24915 ±<br>0.0022 | 0.70848 ±<br>0.0057 | 0.28748 ±<br>0.0067 | 0.25371 ±<br>0.0019 | 0.14421 ±<br>0.0017 | 0.41815 ±<br>0.0090 | 1.00000 ±<br>0.0000 | 0.16910 ±<br>0.0021 | 0.12423 ±<br>0.0018 | 1.00000 ±<br>0.0000 |  |

**Table S6.** Rate of synonymous divergence Ks-JC, detected between different alleles for gene *coxa* of *Wolbachia* infecting *C. parallelus*. SD: Standard deviation, calculated using the formula of the standard deviation of the ratio given by Neter *et al*. (1978) as proposed by Raychoudhury *et al*. (2009).

| Allele x | Allele y | Ks | ±SD | Divergence (years) | ±years |
| --- | --- | --- | --- | --- | --- |
| H1 | H2 | 0.365 | 0.053 | 40.588.889 | 5.872.561 |
| H1 | H3 | 0.363 | 0.053 | 40.377.778 | 5.854.403 |
| H1 | H4 | 0.037 | 0.021 | 4.111.111 | 2.302.144 |
| H1 | H5 | 0.365 | 0.053 | 40.588.889 | 5.872.561 |
| H1 | H6 | 0.346 | 0.052 | 38.433.333 | 5.801.173 |
| H2 | H3 | 0.190 | 0.043 | 21.133.333 | 4.776.953 |
| H2 | H4 | 0.385 | 0.053 | 42.788.889 | 5.934.819 |
| H2 | H5 | 0.012 | 0.012 | 1.344.444 | 1.333.422 |
| H2 | H6 | 0.012 | 0.012 | 1.344.444 | 1.333.422 |
| H3 | H4 | 0.383 | 0.053 | 42.566.667 | 5.917.255 |
| H3 | H5 | 0.175 | 0.042 | 19.433.333 | 4.623.863 |
| H3 | H6 | 0.175 | 0.042 | 19.433.333 | 4.623.863 |
| H4 | H5 | 0.385 | 0.053 | 42.788.889 | 5.934.819 |
| H4 | H6 | 0.365 | 0.053 | 40.588.889 | 5.872.561 |
| H5 | H6 | 0.025 | 0.017 | 2.722.222 | 1.885.450 |

**Table S7.** Rate of synonymous divergence Ks-JC, detected between different alleles for gene *fbpA* of *Wolbachia* infecting *C. parallelus*. SD: Standard deviation, calculated using the formula of the standard deviation of the ratio given by Neter *et al*. (1978) as proposed by Raychoudhury *et al*. (2009).

| Allele x | Allele y | Ks | ±SD | Divergence (years) | ±years |
| --- | --- | --- | --- | --- | --- |
| H1 | H2 | 0.659 | 0.047 | 73.177.778 | 5.199.701 |
| H1 | H2 | 0.040 | 0.019 | 4.444.444 | 2.147.990 |
| H1 | H3 | 0.681 | 0.046 | 75.622.222 | 5.108.707 |
| H1 | H5 | 0.030 | 0.017 | 3.311.111 | 1.863.101 |
| H2 | H2 | 0.661 | 0.047 | 73.455.556 | 5.196.782 |
| H2 | H3 | 0.010 | 0.010 | 1.088.889 | 1.081.110 |
| H2 | H5 | 0.660 | 0.047 | 73.366.667 | 5.197.733 |
| H2 | H3 | 0.683 | 0.046 | 75.922.222 | 5.103.356 |
| H2 | H5 | 0.072 | 0.025 | 7.944.444 | 2.826.638 |
| H3 | H5 | 0.682 | 0.046 | 75.822.222 | 5.104.997 |

**Table S8.** Rate of synonymous divergence Ks-JC, detected between different alleles for gene *gatB* of *Wolbachia* infecting *C. parallelus*. SD: Standard deviation, calculated using the formula of the standard deviation of the ratio given by Neter *et al*. (1978) as proposed by Raychoudhury *et al*. (2009).

| Allele x | Allele y | Ks | ±SD | Divergence (years) | ±years |
| --- | --- | --- | --- | --- | --- |
| H1 | H2 | 0.455 | 0.056 | 50.555.556 | 6.238.165 |
| H1 | H3 | 0.013 | 0.013 | 1.422.222 | 1.409.711 |
| H1 | H4 | 0.506 | 0.056 | 56.244.444 | 6.276.681 |
| H1 | H5 | 0.000 | 0.000 | 0 | 0 |
| H2 | H3 | 0.432 | 0.056 | 48.000.000 | 6.205.388 |
| H2 | H4 | 0.026 | 0.018 | 2.877.778 | 1.991.931 |
| H2 | H5 | 0.455 | 0.056 | 50.555.556 | 6.238.165 |
| H3 | H4 | 0.482 | 0.056 | 53.500.000 | 6.272.865 |
| H3 | H5 | 0.013 | 0.013 | 1.422.222 | 1.409.711 |
| H4 | H5 | 0.506 | 0.056 | 56.244.444 | 6.276.681 |

**Table S9.** Rate of synonymous divergence Ks-JC, detected between different alleles for gene *ftsZ* of *Wolbachia* infecting *C. parallelus*. SD: Standard deviation, calculated using the formula of the standard deviation of the ratio given by Neter *et al*. (1978) as proposed by Raychoudhury *et al*. (2009).

| Allele x | Allele y | Ks | ±SD | Divergence (years) | ±years |
| --- | --- | --- | --- | --- | --- |
| H1 | H2 | 0.765 | 0.042 | 85.022.222 | 4.659.425 |
| H1 | H3 | 0.020 | 0.014 | 2.188.889 | 1.523.871 |
| H1 | H4 | 0.010 | 0.010 | 1.088.889 | 1.079.374 |
| H1 | H5 | 0.739 | 0.043 | 82.055.556 | 4.830.665 |
| H2 | H3 | 0.767 | 0.042 | 85.255.556 | 4.648.771 |
| H2 | H4 | 0.739 | 0.043 | 82.055.556 | 4.830.665 |
| H2 | H5 | 0.030 | 0.017 | 3.355.556 | 1.887.422 |
| H3 | H4 | 0.030 | 0.017 | 3.311.111 | 1.864.552 |
| H3 | H5 | 0.741 | 0.043 | 82.277.778 | 4.822.682 |
| H4 | H5 | 0.713 | 0.045 | 79.188.889 | 4.974.128 |

**Table S10.**
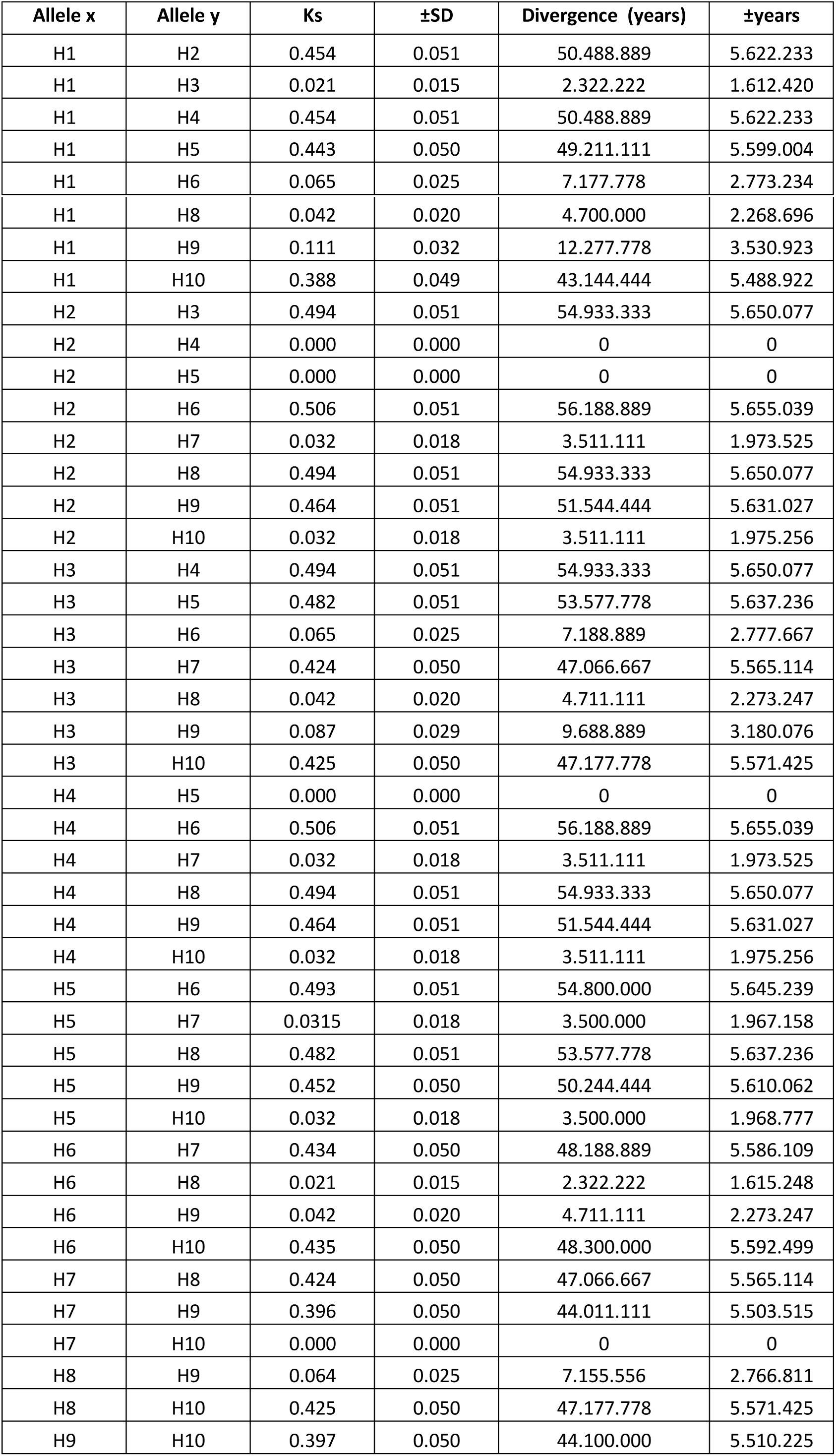
Rate of synonymous divergence Ks-JC, detected between different alleles for gene *hcpA* of *Wolbachia* infecting *C. parallelus*. SD: Standard deviation, calculated using the formula of the standard deviation of the ratio given by Neter *et al*. (1978) as proposed by Raychoudhury *et al*. (2009).

**Table S11.** Synonymous divergence Ks-JC between the main ST belonging to supergroup F detected in both subspecies of *C. parallelus*. SD: Standard deviation, calculated using the formula of the standard deviation of the ratio given by Neter *et al*. (1978) as proposed by Raychoudhury *et al*. (2009).

| Allele x | Allele y | Ks | ±SD | Divergence (years) | ±years |
| --- | --- | --- | --- | --- | --- |
| ST10 | ST11 | 0.0109 | 0.005 | 1.211.111 | 535.687 |
| ST10 | ST24 | 0.0376 | 0.009 | 4.177.778 | 982.023 |
| ST10 | ST25 | 0.0241 | 0.007 | 2.677.778 | 791.342 |
| ST11 | ST24 | 0.0445 | 0.010 | 4.944.444 | 1 064.878 |
| ST11 | ST25 | 0.0308 | 0.008 | 3.422.222 | 891.855 |
| ST24 | ST25 | 0.0131 | 0.005 | 1.455.556 | 587.295 |

**Table S12.** Synonymous divergence Ks-JC between the main ST belonging to supergroup B detected in both subspecies of *C. parallelus*. SD: Standard deviation, calculated using the formula of the standard deviation of the ratio given by Neter *et al*. (1978) as proposed by Raychoudhury *et al*. (2009).

| Allele x | Allele y | Ks | ±SD | Divergence (years) | ±years |
| --- | --- | --- | --- | --- | --- |
| ST13 | ST31 | 0.014 | 0.006 | 1.566.667 | 633.171 |
| ST6 | ST31 | 0.012 | 0.005 | 1.311.111 | 579.907 |
| ST13 | ST33 | 0.012 | 0.005 | 1.311.111 | 580.524 |
| ST14 | ST31 | 0.012 | 0.005 | 1.311.111 | 579.907 |
| ST17 | ST31 | 0.012 | 0.005 | 1.311.111 | 579.677 |
| ST3 | ST31 | 0.009 | 0.005 | 1.044.444 | 518.213 |
| ST6 | ST33 | 0.009 | 0.005 | 1.044.444 | 518.764 |
| ST7 | ST31 | 0.009 | 0.005 | 1.044.444 | 518.007 |
| ST14 | ST33 | 0.009 | 0.005 | 1.044.444 | 518.764 |
| ST16 | ST31 | 0.009 | 0.005 | 1.044.444 | 518.007 |
| ST17 | ST33 | 0.009 | 0.005 | 1.044.444 | 518.564 |
| ST3 | ST33 | 0.007 | 0.004 | 788.889 | 451.377 |
| ST7 | ST33 | 0.007 | 0.004 | 788.889 | 451.202 |
| ST13 | ST32 | 0.007 | 0.004 | 788.889 | 451.292 |
| ST16 | ST33 | 0.007 | 0.004 | 788.889 | 451.202 |
| ST6 | ST16 | 0.007 | 0.004 | 777.778 | 447.947 |
| ST32 | ST31 | 0.007 | 0.004 | 777.778 | 447.733 |
| ST3 | ST13 | 0.005 | 0.003 | 522.222 | 367.622 |
| ST3 | ST16 | 0.005 | 0.003 | 522.222 | 367.476 |
| ST6 | ST14 | 0.005 | 0.003 | 522.222 | 367.622 |
| ST6 | ST17 | 0.005 | 0.003 | 522.222 | 367.476 |
| ST6 | ST32 | 0.005 | 0.003 | 522.222 | 367.622 |
| ST7 | ST13 | 0.005 | 0.003 | 522.222 | 367.476 |
| ST7 | ST16 | 0.005 | 0.003 | 522.222 | 367.334 |
| ST13 | ST16 | 0.005 | 0.003 | 522.222 | 367.476 |
| ST14 | ST32 | 0.005 | 0.003 | 522.222 | 367.622 |
| ST17 | ST32 | 0.005 | 0.003 | 522.222 | 367.476 |
| ST32 | ST33 | 0.005 | 0.003 | 522.222 | 367.691 |
| ST3 | ST6 | 0.002 | 0.002 | 255.556 | 257.478 |
| ST3 | ST14 | 0.002 | 0.002 | 255.556 | 257.478 |
| ST3 | ST17 | 0.002 | 0.002 | 255.556 | 257.375 |
| ST3 | ST32 | 0.002 | 0.002 | 255.556 | 257.478 |
| ST6 | ST7 | 0.002 | 0.002 | 255.556 | 257.375 |
| ST6 | ST13 | 0.002 | 0.002 | 255.556 | 257.478 |
| ST7 | ST14 | 0.002 | 0.002 | 255.556 | 257.375 |
| ST7 | ST17 | 0.002 | 0.002 | 255.556 | 257.276 |
| ST7 | ST32 | 0.002 | 0.002 | 255.556 | 257.375 |
| ST13 | ST14 | 0.002 | 0.002 | 255.556 | 257.478 |
| ST13 | ST17 | 0.002 | 0.002 | 255.556 | 257.375 |
| ST14 | ST16 | 0.002 | 0.002 | 255.556 | 257.375 |
| ST16 | ST17 | 0.002 | 0.002 | 255.556 | 257.276 |
| ST16 | ST32 | 0.002 | 0.002 | 255.556 | 257.375 |
| ST33 | ST31 | 0.002 | 0.002 | 255.556 | 257.300 |
| ST3 | ST7 | 0.000 | 0.000 | 0 | 0 |
| ST14 | ST17 | 0.000 | 0.000 | 0 | 0 |

**Table 13s.**
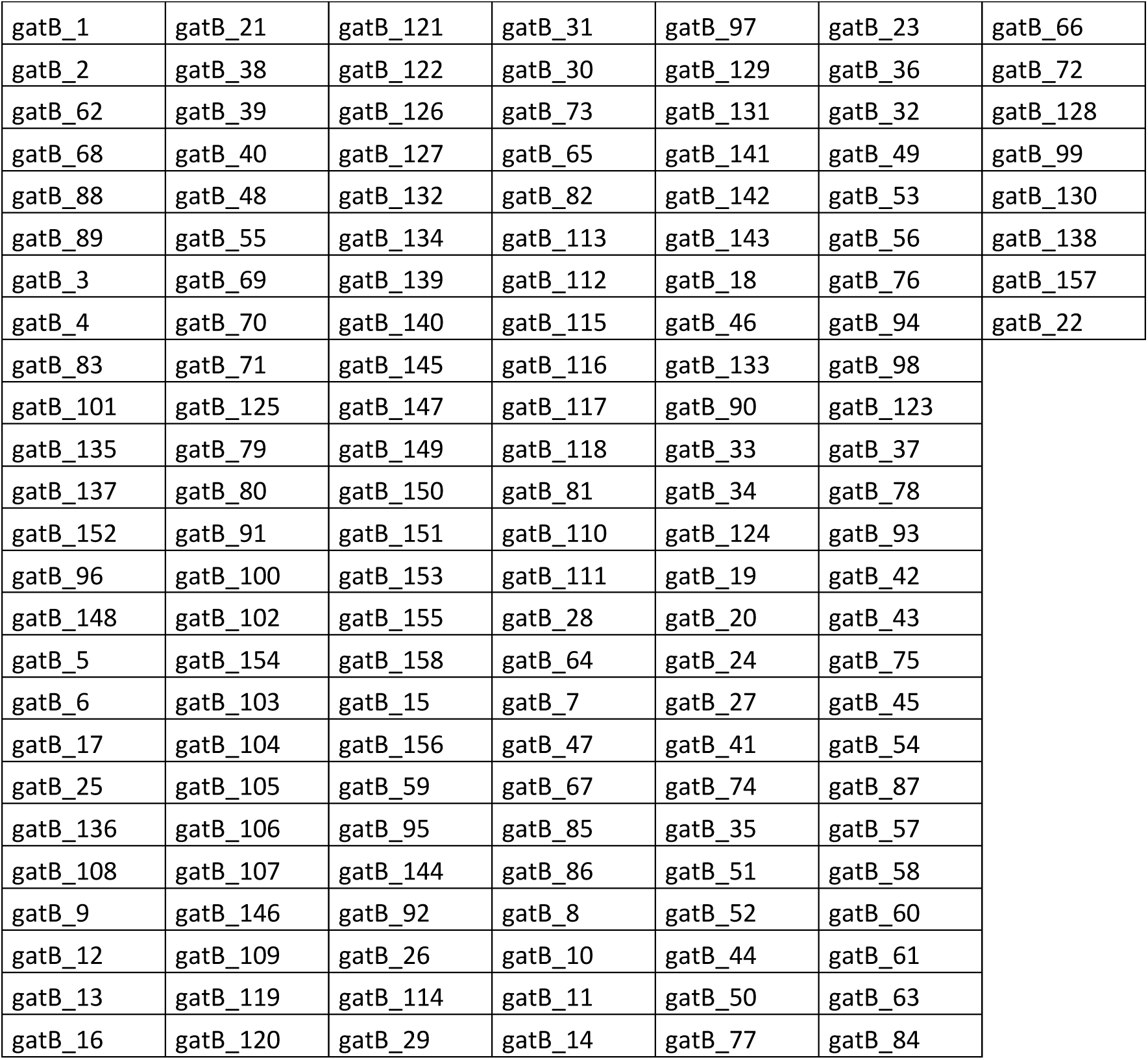
Alleles includes in phylogenetic analysis of gatB

**Table 14s.**
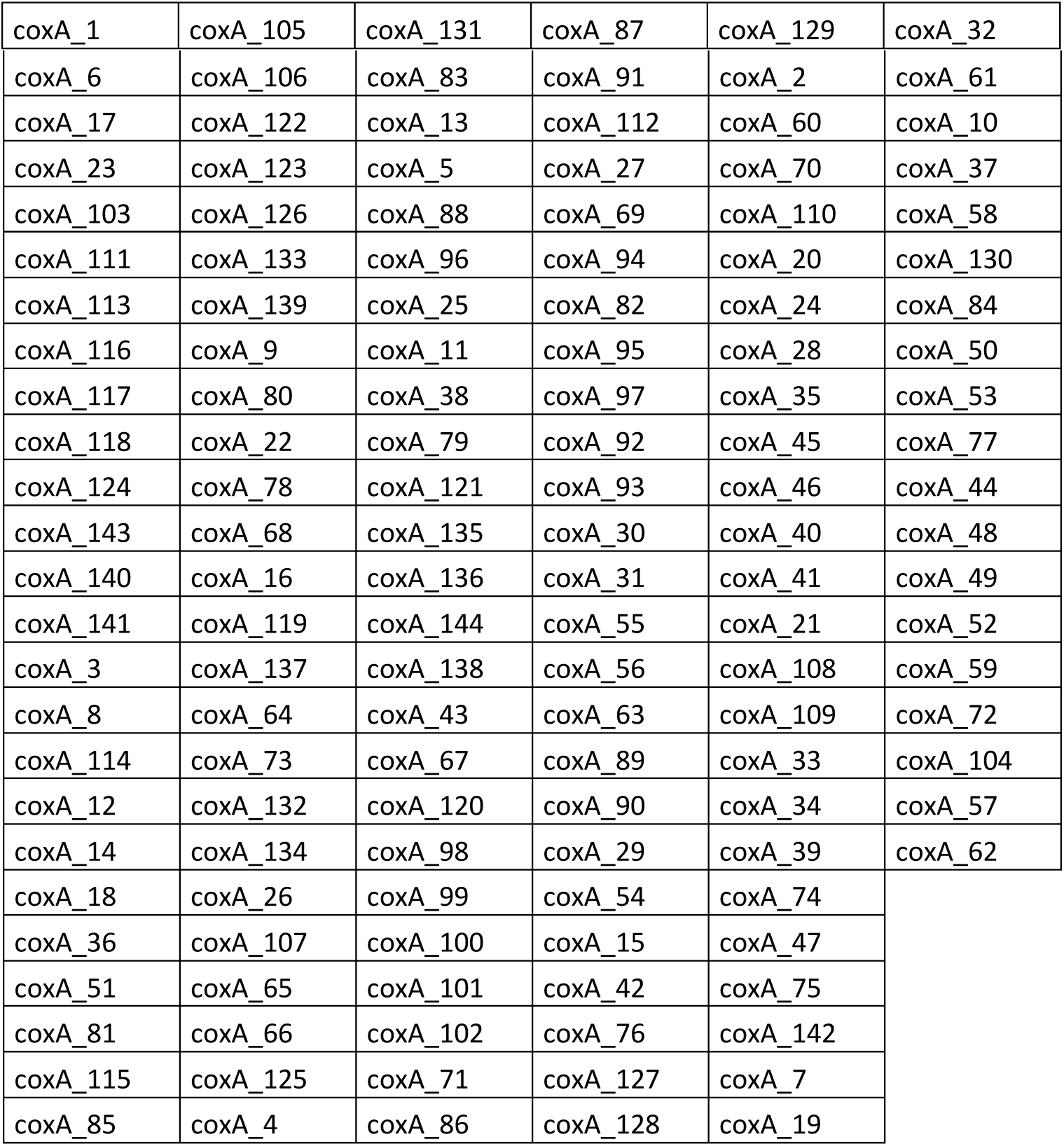
Alleles includes in phylogenetic analysis of coxA

**Table 15s.**
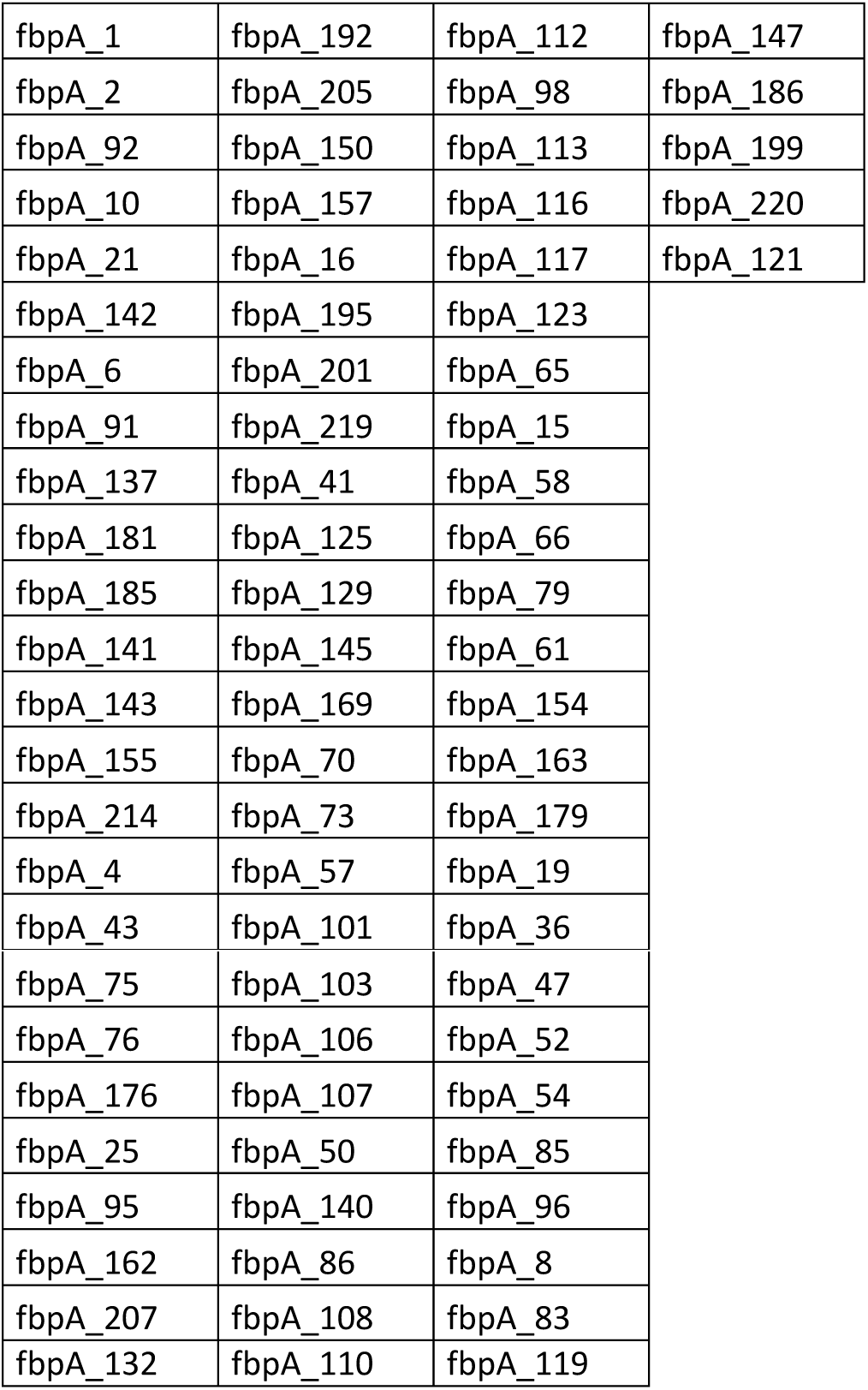
Alleles includes in phylogenetic analysis of fbpA

**Table 16s.**
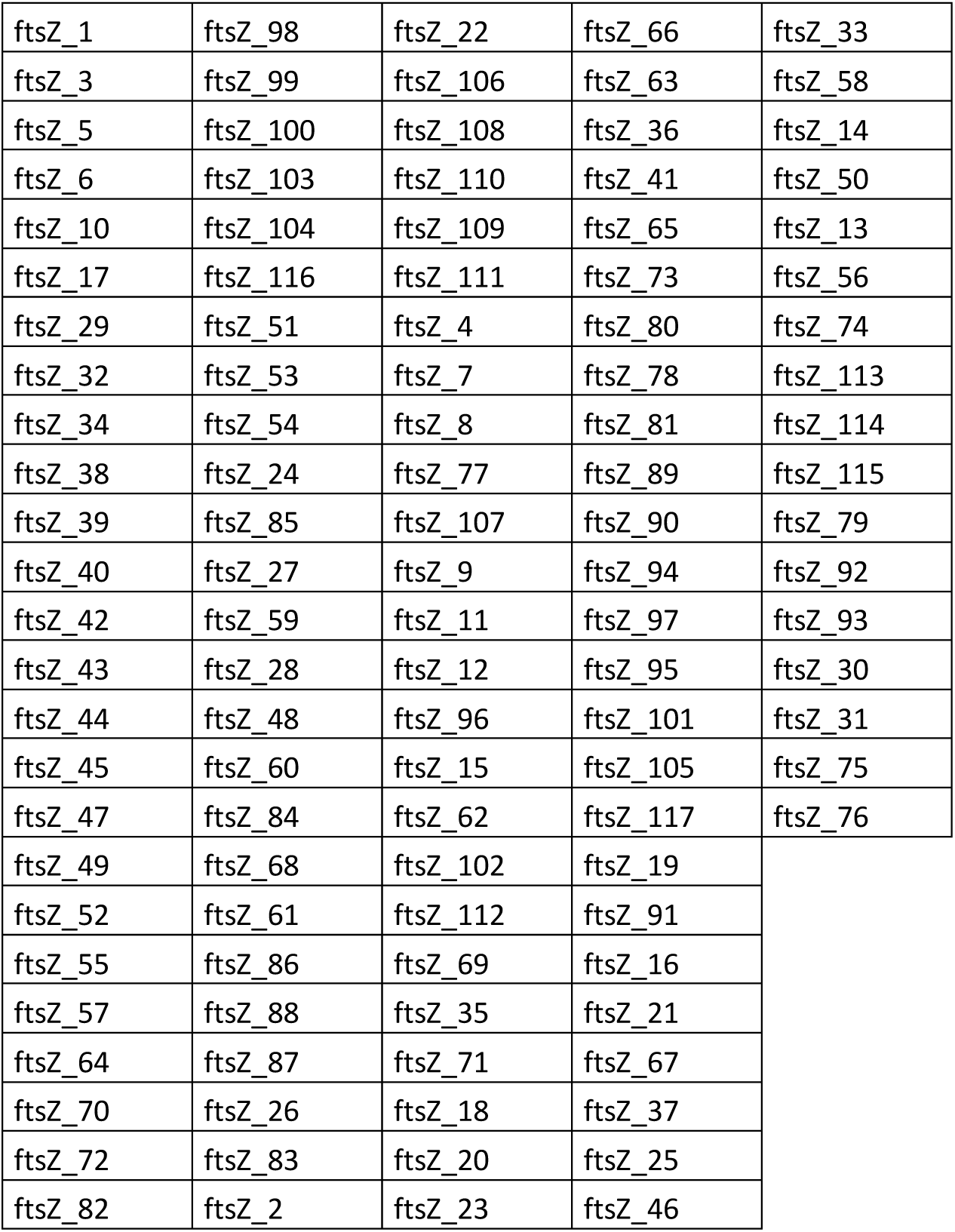
Alleles includes in phylogenetic analysis of ftsZ

**Table 17s.**
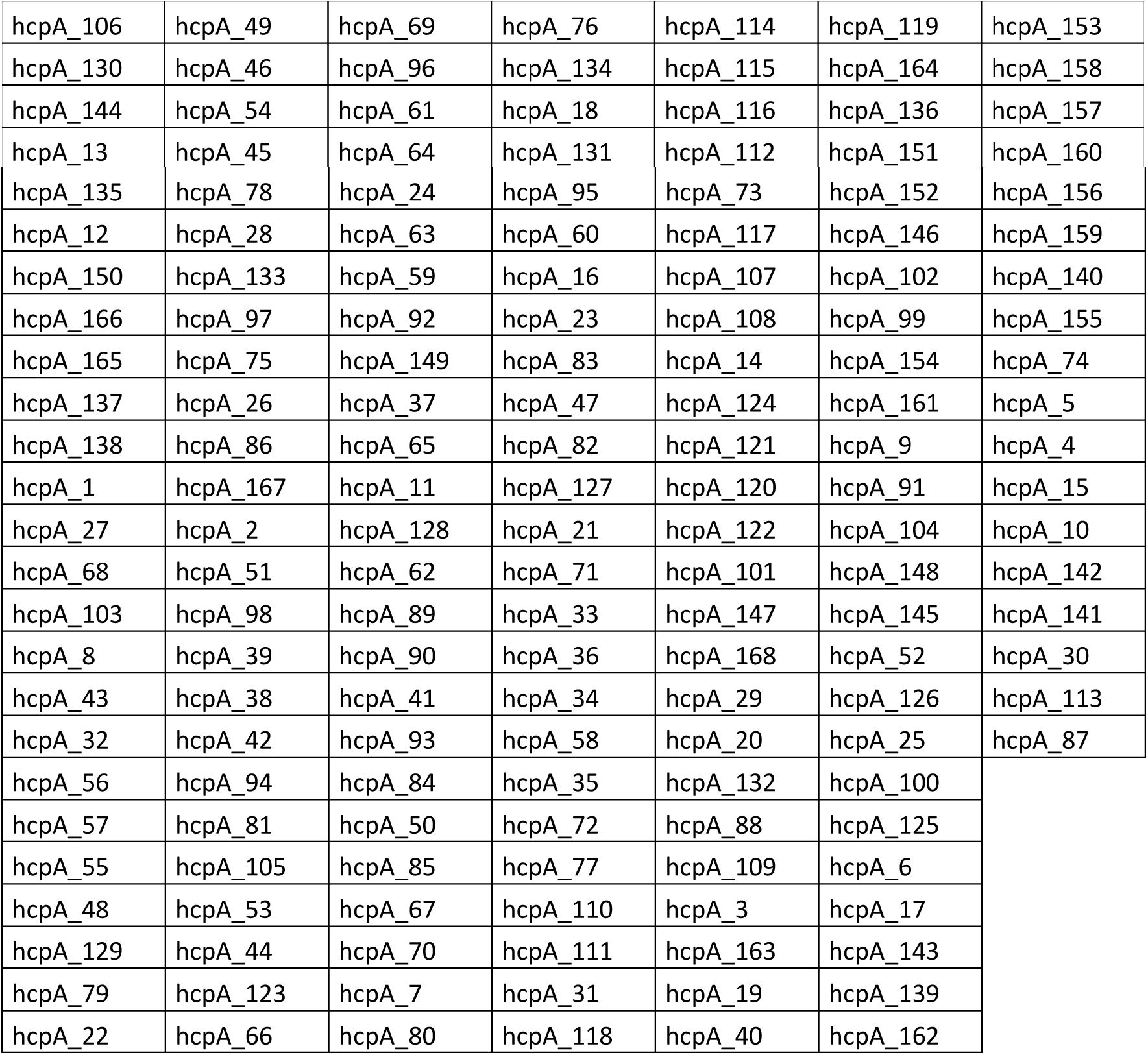
Alleles includes in phylogenetic analysis of hcpA

**Table 18s.**
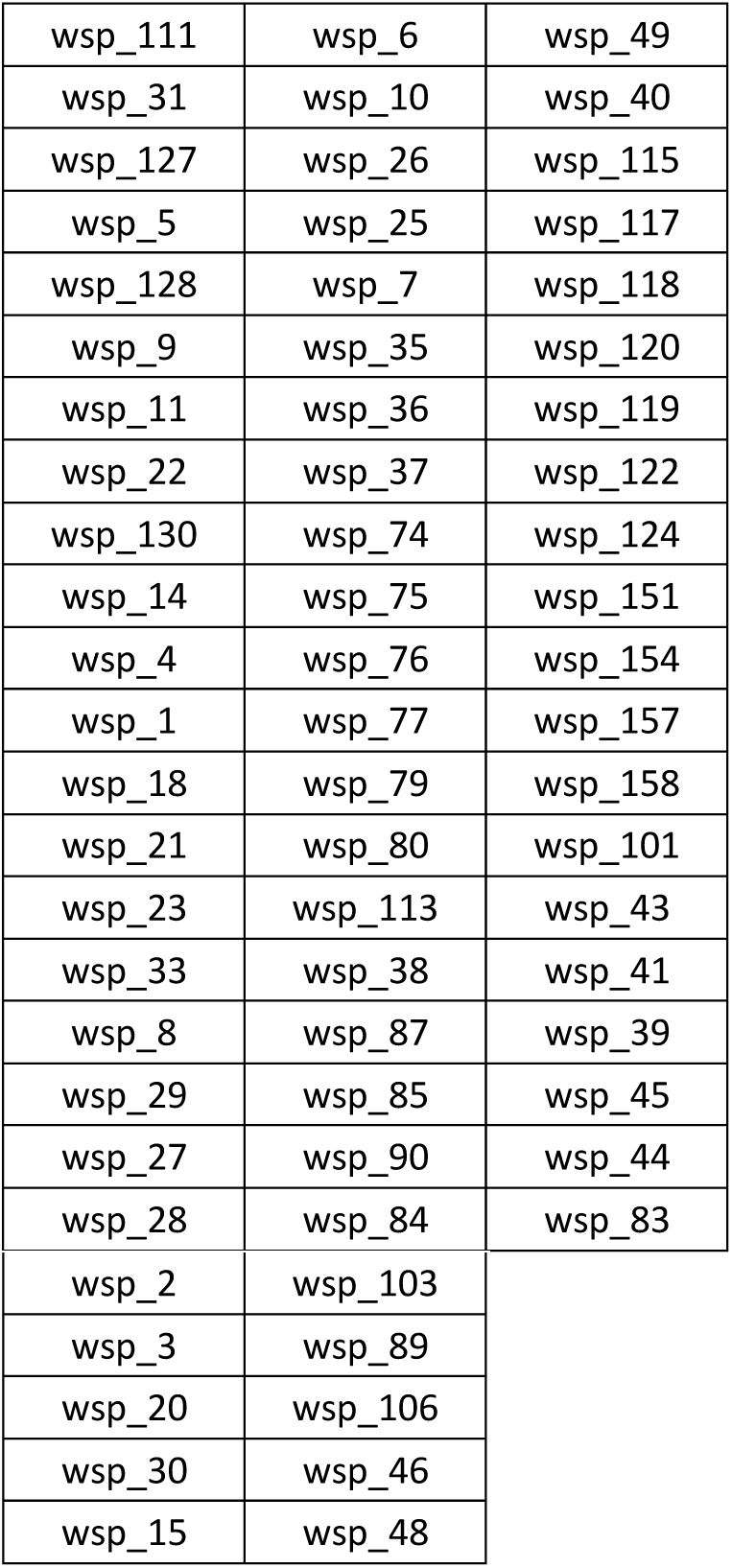
Alleles includes in phylogenetic analysis of wsp

## Supplemental figures

**Fig S1.**
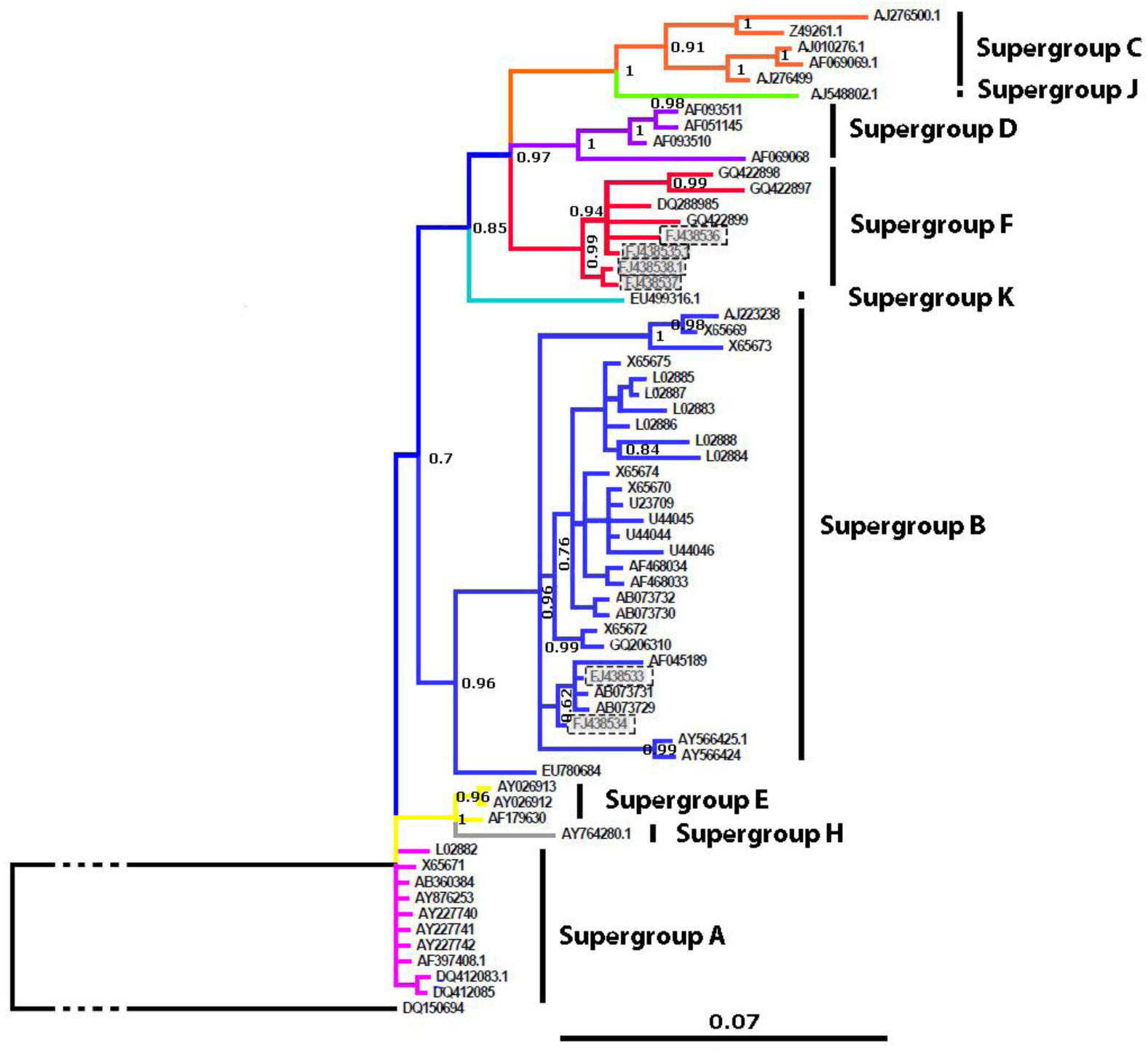
(Online colour figure) Bayesian phylogenetic tree based on *16S rRNA* gene. Outgroup: *E. coli. Wolbachia* infection in *C. parallelus* is shaded is framed grey.

**Fig S2.**
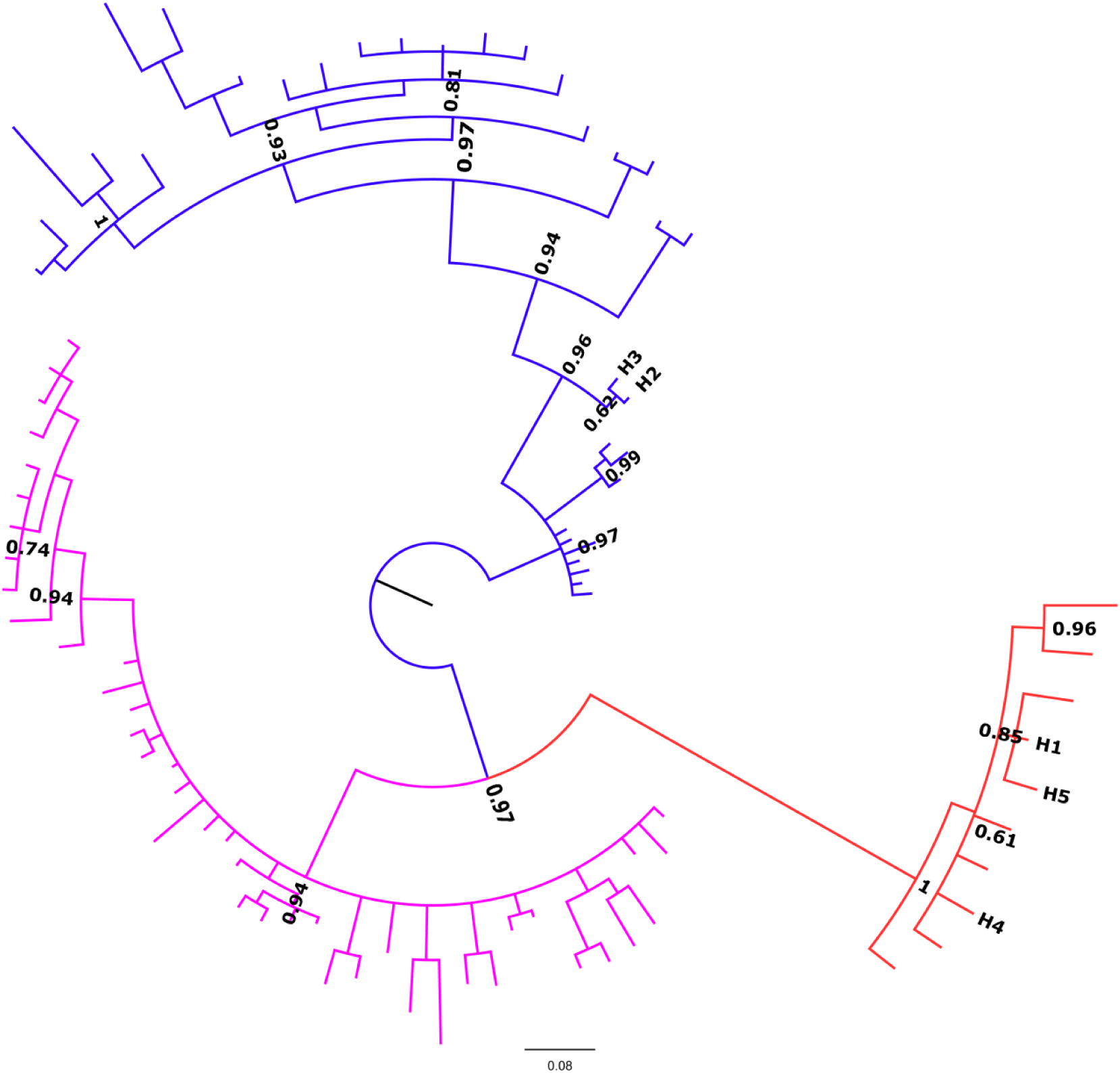
Unrooted phylogenetic tree of *fbpA*, obtained by Bayesian inference. Alleles described in *C. parallelus* appear named H1 to H5. Posterior probabilities are shown in the nodes. The color code encodes the supergroup A (pink), B (blue), D (green), F (red) and H (purple). Posterior probabilities are shown in the nodes.

**Fig S3.**
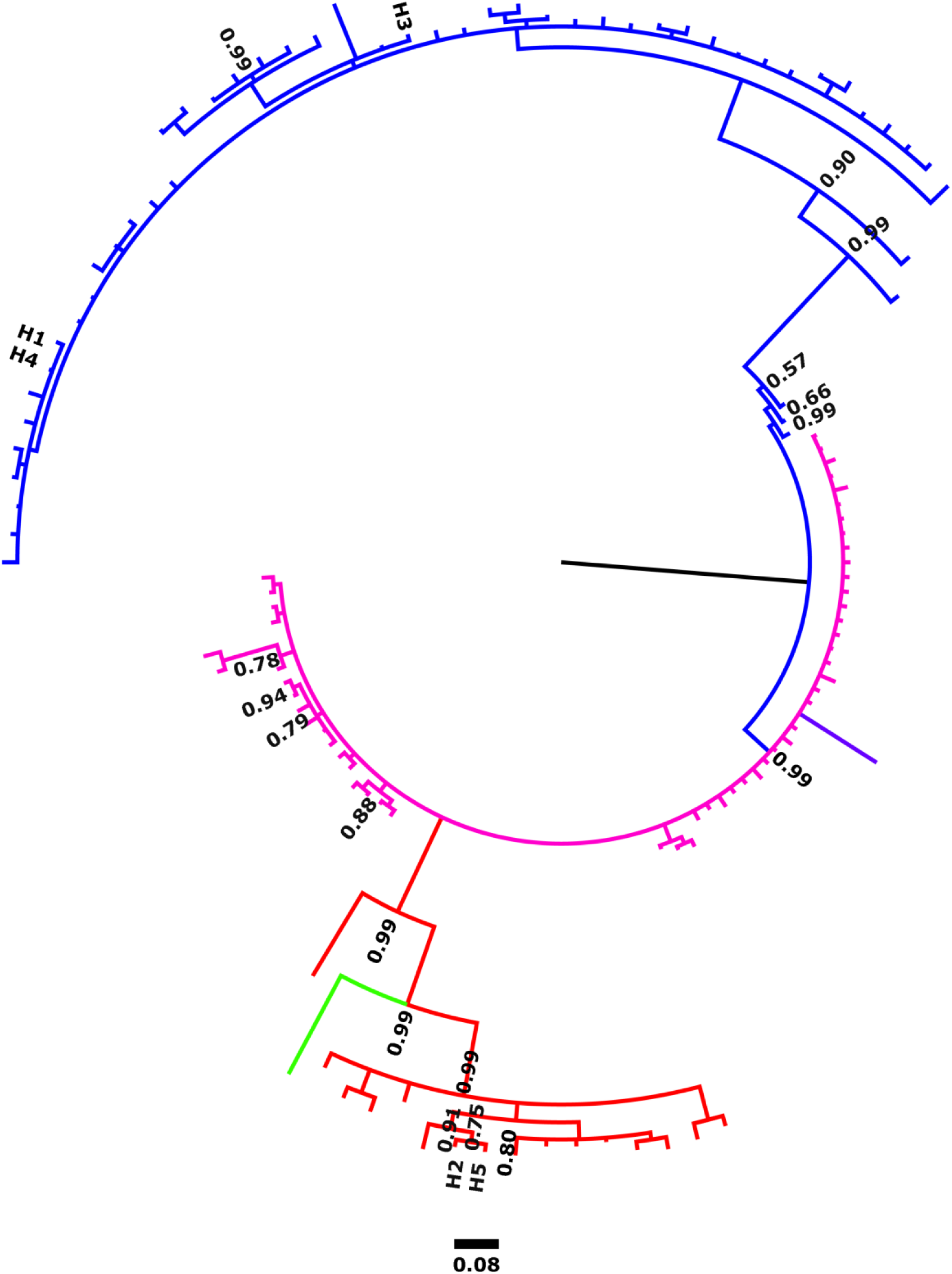
Unrooted phylogenetic tree of *ftsZ*, obtained by Bayesian inference. Alleles described in *C. parallelus* appear named H1 to H5. Posterior probabilities are shown in the nodes. The color code encodes the supergroup A (pink), B (blue), D (green), F (red) and H (purple). Posterior probabilities are shown in the nodes.

**Fig S4.**
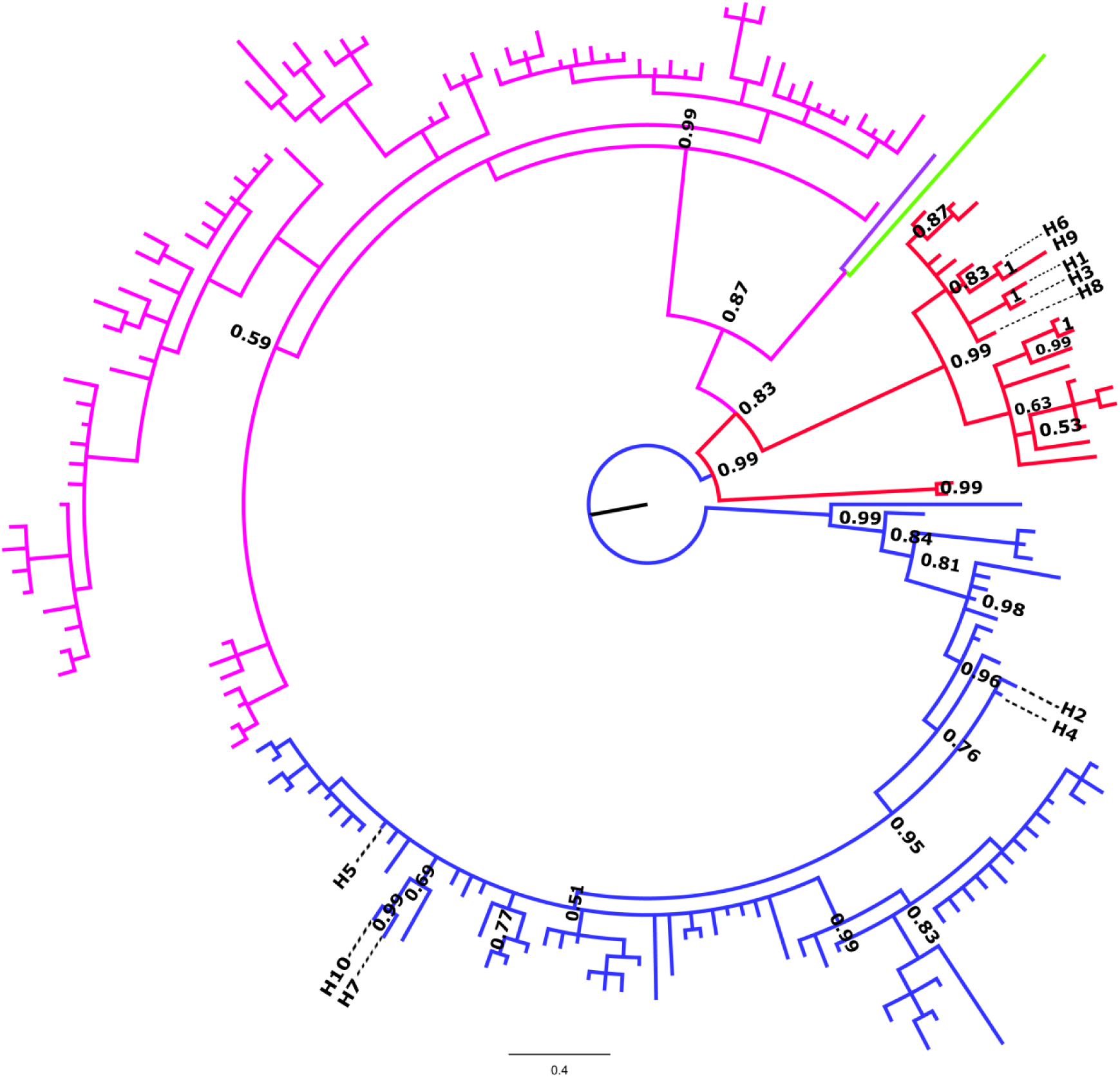
Unrooted phylogenetic tree of *hcpA*, obtained by Bayesian inference. Alleles described in *C. parallelus* appear named H1 to H10. Posterior probabilities are shown in the nodes. The color code encodes the supergroup A (pink), B (blue), D (green), F (red) and H (purple). Posterior probabilities are shown in the nodes.

**Fig S5.**
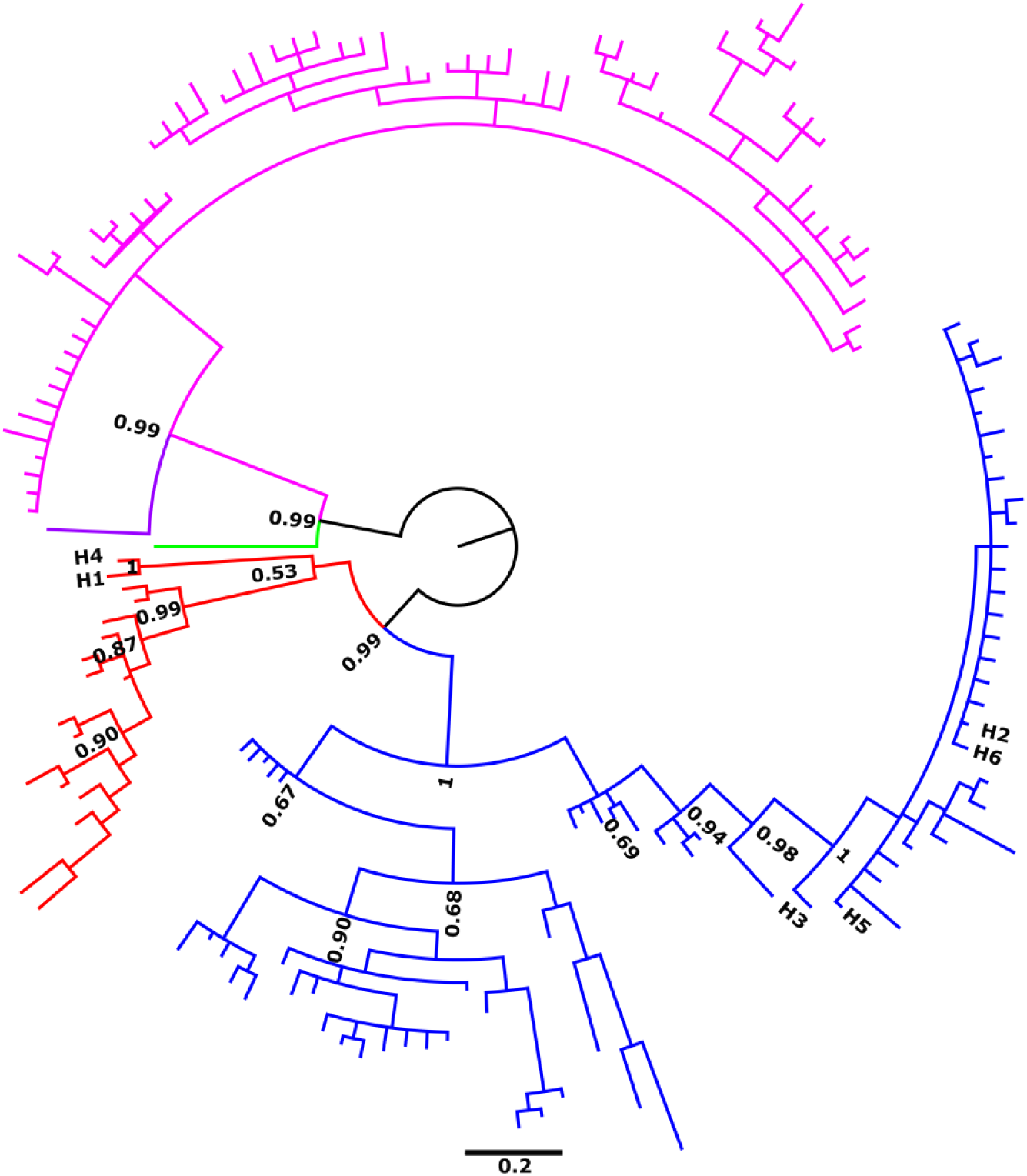
Unrooted phylogenetic tree of *coxA*, obtained by Bayesian inference. Alleles described in *C. parallelus* appear named H1 to H6. Posterior probabilities are shown in the nodes. The color code encodes the supergroup A (pink), B (blue), D (green), F (red) and H (purple). Posterior probabilities are shown in the nodes.

**Fig S6.**
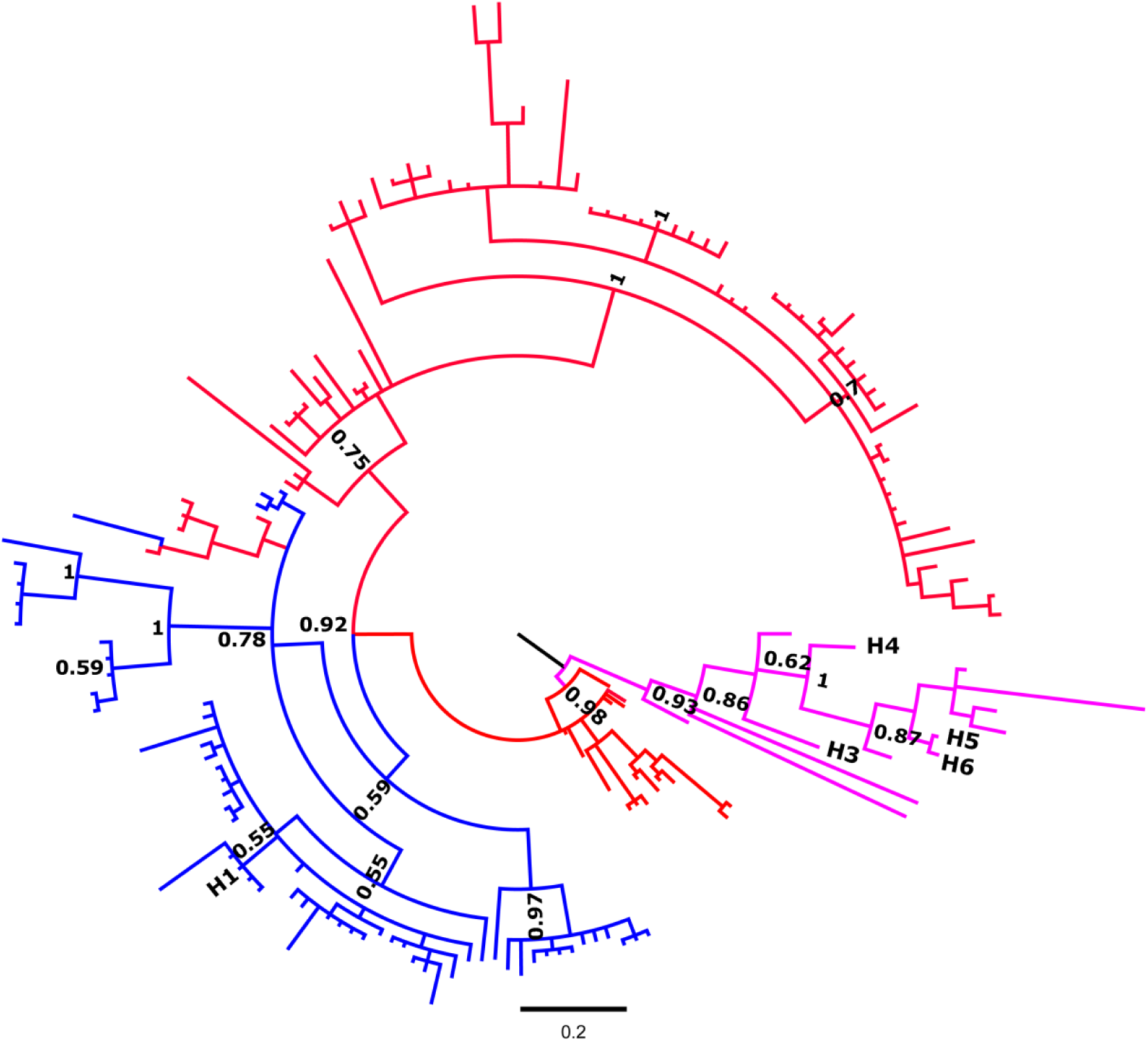
Unrooted phylogenetic tree of *wsp*, obtained by Bayesian inference. Alleles described in *C. parallelus* appear named H1 to H16. Posterior probabilities are shown in the nodes. The color code encodes the supergroup A (pink), B (blue), D (green), F (red) and H (purple). Posterior probabilities are shown in the nodes.

**Fig S7.**
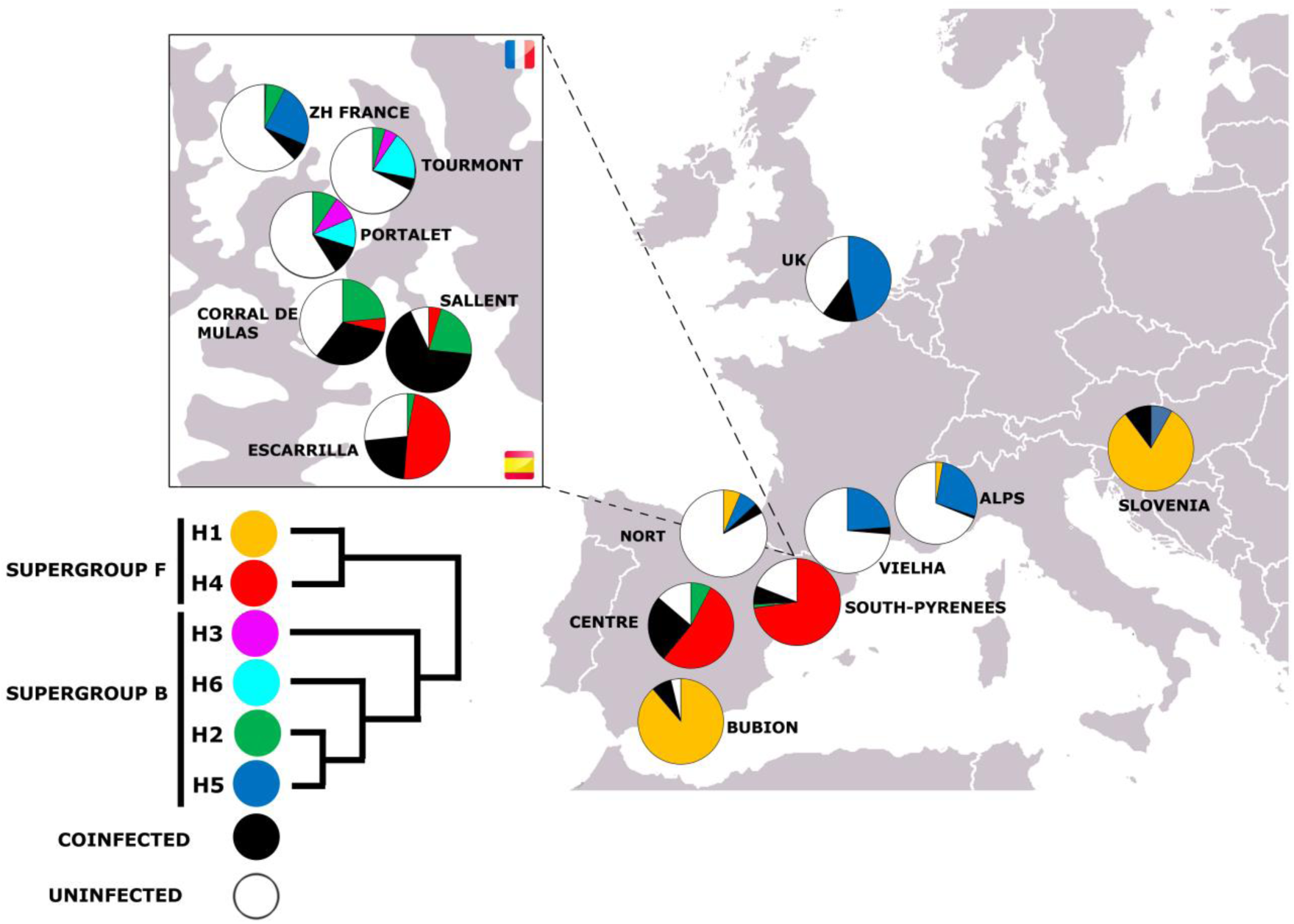
Geographical distribution of alleles detected for gene *coxA*

**Fig S8.**
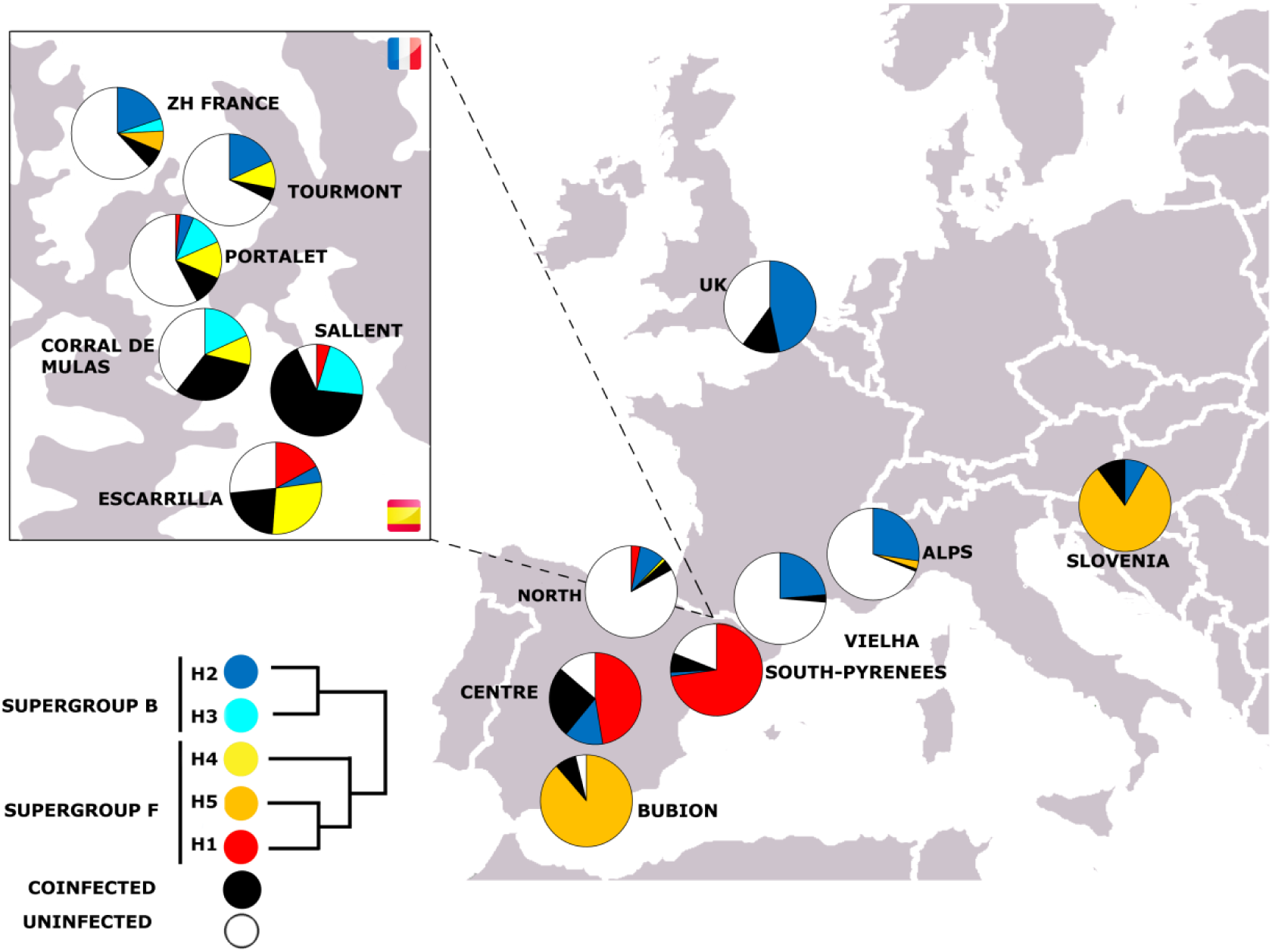
Geographical distribution of alleles detected for gene *fbpA*

**Fig S9.**
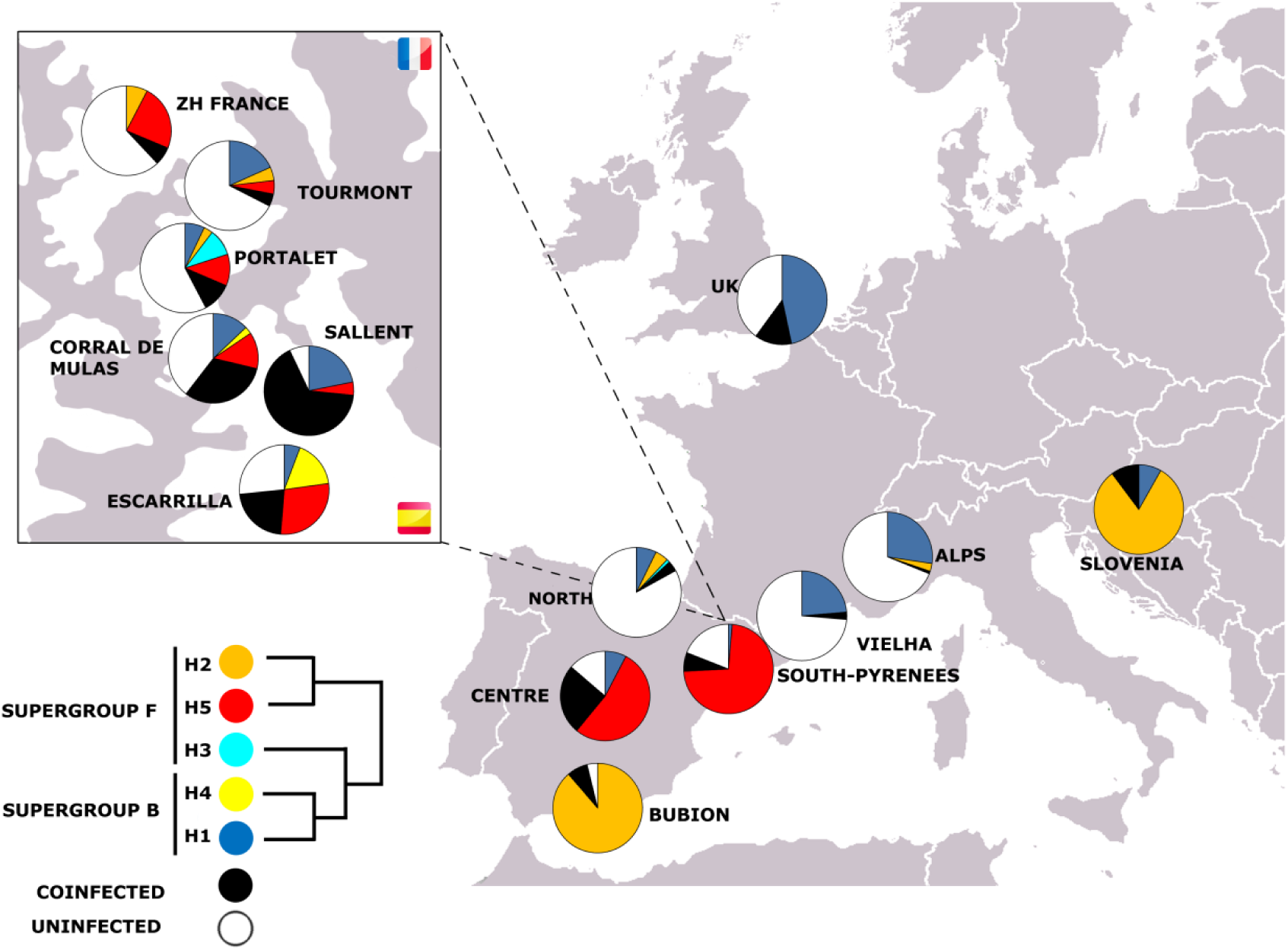
Geographical distribution of alleles detected for gene *ftsZ*

**Fig S10.**
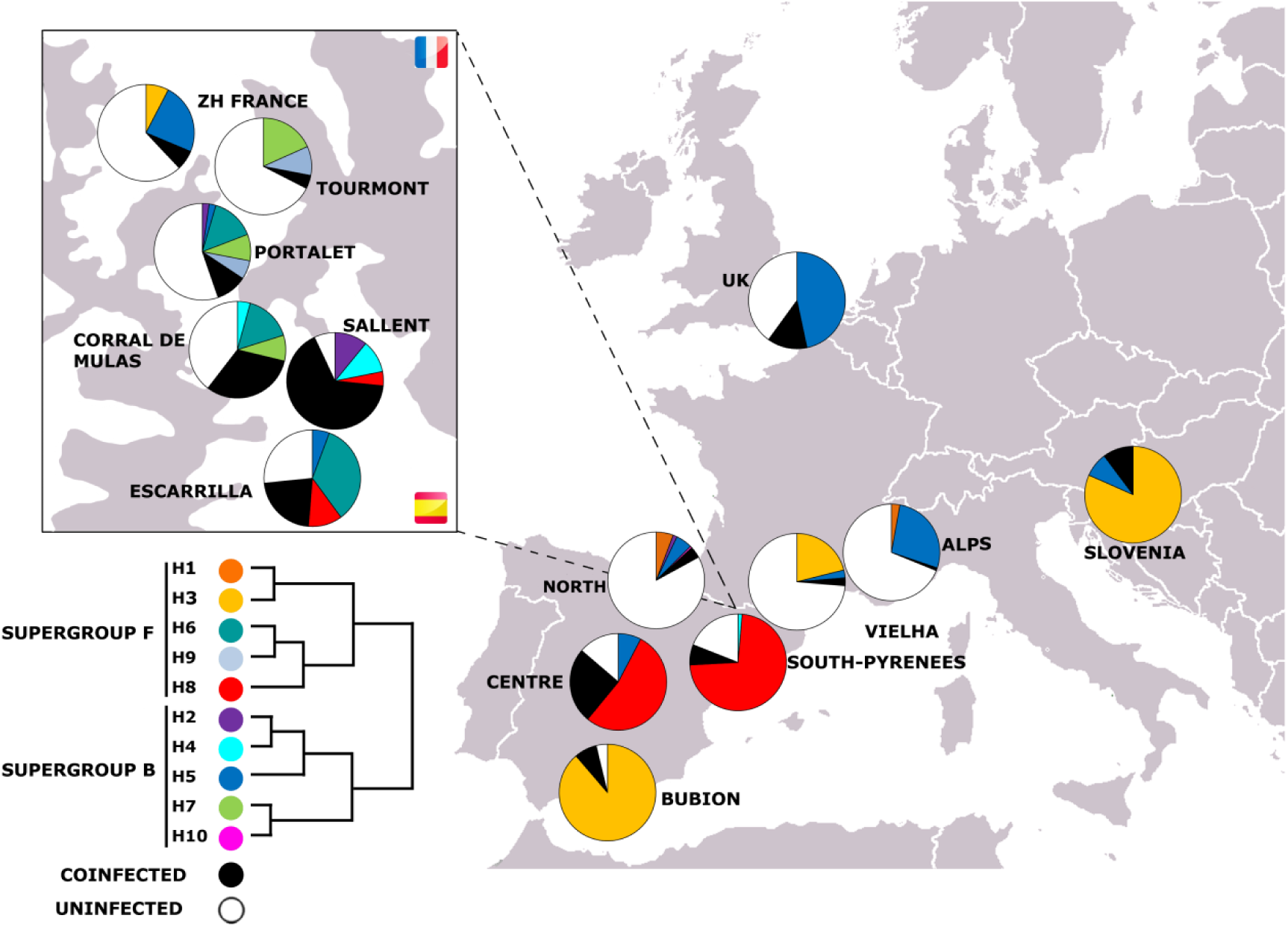
Geographical distribution of alleles detected for gene *hcpA*

**Fig S11.**
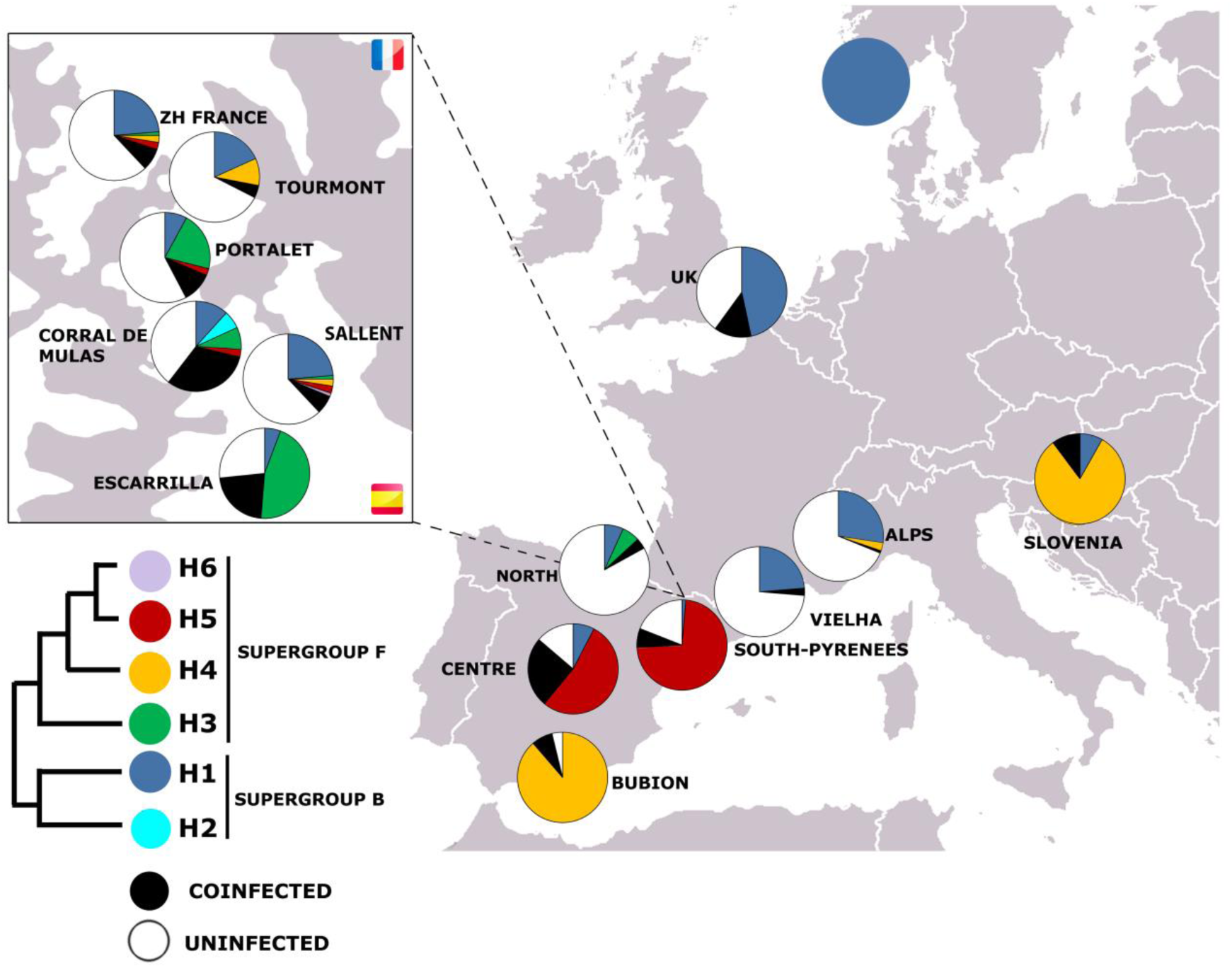
Geographical distribution of alleles detected for gene *wsp*

**Fig S12.**
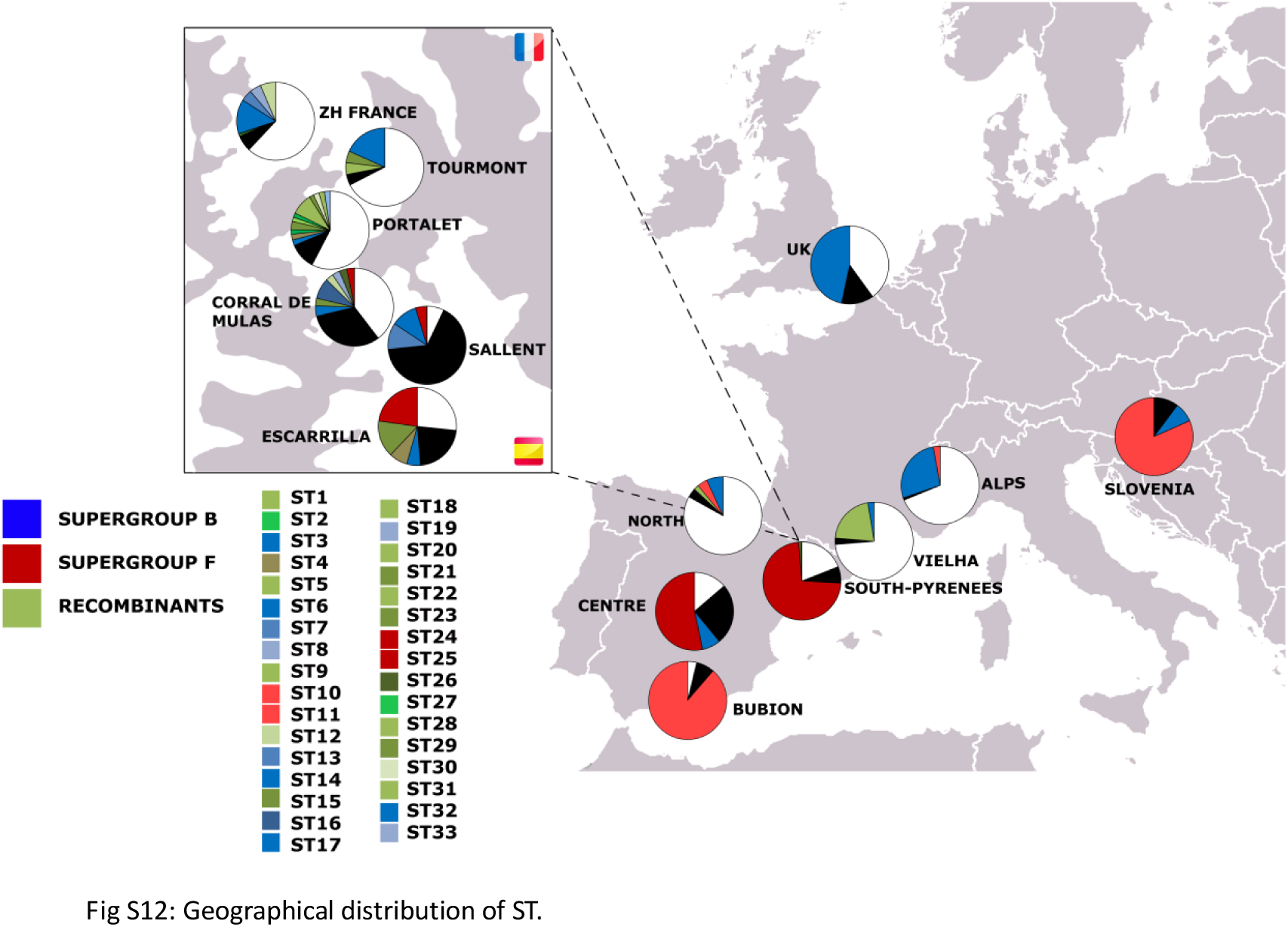
Geographical distribution of ST.

